# Metacommunity dynamics and the detection of species associations in co-occurrence analyses: why patch disturbance matters

**DOI:** 10.1101/2021.06.14.448327

**Authors:** Vincent Calcagno, Nik J. Cunniffe, Frédéric M. Hamelin

## Abstract

- 1: Many statistical methods attempt to detect species associations – and so infer inter-specific interactions – from species co-occurrence patterns. Habitat heterogeneity and out-of-equilibrium colonization histories are well recognized as potentially causing species associations, even when interactions are absent. The potential for patch disturbance, a classical component of metacommunity dynamics, to also drive spurious species associations has however been over-looked.
- 2: Using a new general metacommunity model, we derive mathematical predictions regarding how patch disturbance would affect the patterns of species associations detected in “null” co-occurrence matrices. We also conduct numerical simulations to test our predictions and to compare the performance of several widespread statistical methods, including direct tests of pairwise independence, matrix permutation approaches and joint species distribution modelling.
- 3: We show how classical metacommunity dynamics can produce statistical associations, both positive and negative, even when species do not interact, when there is no habitat heterogeneity, and at equilibrium. This occurs as soon as there is some rate of patch disturbance (i.e. simultaneous extinction of several species in a patch) and/or a finite life-span of patches, a common feature of a broad range of plant, animal or microbial systems.
- 4: Patch disturbance can compromise species co-occurrence analyses and cause the artefactual detection of species associations if not taken into account. Including patch age (i.e. the time since the last patch disturbance event) as a covariate in a joint species distribution model can resolve the artefact. However, this requires additional data that often are not available in practice. We argue that the consequences of patch disturbance should not be underestimated when analysing species distribution patterns in metacommunity-like systems.

## 1 Introduction

Community ecology has long been concerned with inferring interactions between species (e.g. competition or facilitation) from co-occurrence data (Forbes, 1907; Cohen, 1970; Diamond, 1975; Caswell, 1976; Connor and Simberloff, 1979). Recently, metagenomics approaches have renewed this use of co-occurrence patterns as a means to infer species interaction networks, in environments ranging from soils to the human microbiome (Barberán et al., 2012; Faust and Raes, 2012). If species do not interact, intuition suggests the proportion of patches (or hosts) where species co-occur should be the product of the proportions of patches occupied by each species. Testing for pairwise independence, i.e. looking for statistical patterns of association, consequently underpins most methods for inferring species interactions from presence-absence data (Gotelli, 2000; Gotelli and Ulrich, 2012; Ovaskainen and Abrego, 2020). Positive associations (excess of co-occurrences) may indicate facilitation whereas negative associations (deficit of co-occurrences) may indicate competition.

However, it is well known that several pitfalls can compromise this approach, since correlation is not equivalent to interaction (Barner et al., 2018; Blanchet et al., 2020; Molina and Stone, 2020). As a simple example, patch heterogeneity and differential habitat preferences among species (i.e. environmental filtering) can cause species associations, even in the absence of any interaction between the pairs of species involved. For instance, if some patch types host more species than others because of their larger size or more favorable conditions, the occurrence of a species in a patch increases the odds of occurrence of other species, simply because it makes it more likely the patch is of higher quality. This would generate positive association signals if not controlled for. Succession or non-equilibrium dynamics, i.e. species expanding through space, concomitantly or differentially, are also expected to cause similar patterns (D’Amen et al., 2018). More recent statistical and logical frameworks incorporate and test for alternative mechanisms to species interactions (Blois et al., 2014; Ulrich et al., 2017; D’Amen et al., 2018). However, it is commonly expected than in the absence of any of the above confounding factors and others (such as dispersal limitation), or after having appropriately controlled for them, classical extinction/recolonization dynamics (metacommunity dynamics; Leibold et al., 2004) would not introduce particular species associations, unless ecological interactions are actually occurring (Opedal et al., 2020b).

Here we show that a specific component of metacommunity dynamics, patch disturbance, has consequences that have been overlooked so far. Patch disturbance here means the stochastic occurrence of patch extinctions (Hastings, 1980, 2003). Patch disturbance events denote extreme but possibly frequent events, such as fires, droughts, floods and others, causing the local extinction of all, or perhaps some subset of all, species from a patch (Sousa, 1984; Leibold et al., 2004). This can also correspond to the actual destruction and disappearance of a patch, for instance in ecosystems managed by humans, such as agricultural fields or forest stands (Ovaskainen et al., 2017), or in systems for which “patches” are actually hosts with a finite lifespan, such as parasite or microbiote communities (Hamelin et al., 2019; Friedman and Alm, 2012). Patch disturbances are ubiquitous in natural and anthropogenic systems (Pickett and White, 2013; Jentsch and White, 2019). Although naturally regarded as a part of metapopulation and metacommunity frameworks (e.g. Levin and Paine, 1974; Hastings, 1980, 2003; Leibold et al., 2004; Calcagno et al., 2011), patch disturbance is often omitted (e.g. Slatkin, 1974) or confounded with species-specific extinctions (e.g. Thompson et al., 2020). Although this may seem a benine mathematical simplification, we will show here that this omission can have non trivial implications.

We first introduce a general metacommunity model of patch dynamics, that allows for patch disturbance to occur at some, potentially time-varying, rate, and in which patches can have a finite lifespan. This model generalizes many existing particular models. The model is “null” in the sense that species are not interacting, the habitat is completely homogeneous, and all species disperse uniformly with no patch preferences. We analyze the equilibrium occupancies and co-occurrence patterns, from which we generate sample co-occurrence matrices. We then test different often-used approaches to detect species associations on these “null” synthetic data. The methods considered include direct statistical tests of independence (e.g. Veech, 2013), different permutation-based matrix analyses (e.g. Gotelli, 2000), and joint species distribution modelling (e.g. Ovaskainen and Abrego, 2020). In each case, we first derive mathematical predictions concerning the expected performance of the method in question, and then test these predictions with numerical simulations.

We show that the occurrence of patch disturbance will yield characteristic patterns of spurious species associations, for all methods that do not explicitly model the effect of patch-age (time since last disturbance event) as a covariate. We show that a quantity we term species “fastness”, that captures how quickly species occurrence probability recovers after disturbance under the effect of recolonization/extinction dynamics, is a key determinant of the expected patterns of spurious associations. We show that the use of joint species distribution modelling with patch age as an environmental covariate (e.g. Ovaskainen et al., 2017) is the only way to achieve satisfactory performance (absence of spurious signals). Since patch age would often be unknown, we explore whether patch richness (i.e. the number of species in a patch) could suffice as a proxy for patch age, but we find it does not. We suggest the implications of patch disturbance as a constituent of metacommunity dynamics should not be underestimated, and that explicit data on patch-age would generally be required for accurate species co-occurrence analysis in metacommunity-like systems.

## 2 Materials & Methods

### 2.1 A general metacommunity model

We consider a metacommunity model describing the extinction-recolonization dynamics of *s* non-interacting species over a large number of identical patches (Caswell, 1976). Each species (*i*) has its own colonization rate *c_i_* per occupied patch, as well as a constant rate of immigration from outside the metacommunity, *m_i_* (Hastings, 1987). Any species can go extinct in a patch, at some species-dependent rate *e_i_*. In addition, patches can undergo catastrophic disturbances, after which all species in the patch immediately go extinct, irrespective of species composition (Hastings, 1980; Leibold et al., 2004; Calcagno et al., 2011). Such catastrophes occur at rate *μ_x_*, where *x* is the “age” of a patch, i.e. the time since its appearance or since it experienced the last catastrophic disturbance event (Levin and Paine, 1974; Hastings, 2003). Finally, there can be some maximal patch age *X* after which a catastrophic disturbance systematically occurs (Olivieri et al., 1995). The model is illustrated in Fig. 1.

**Figure 1:**
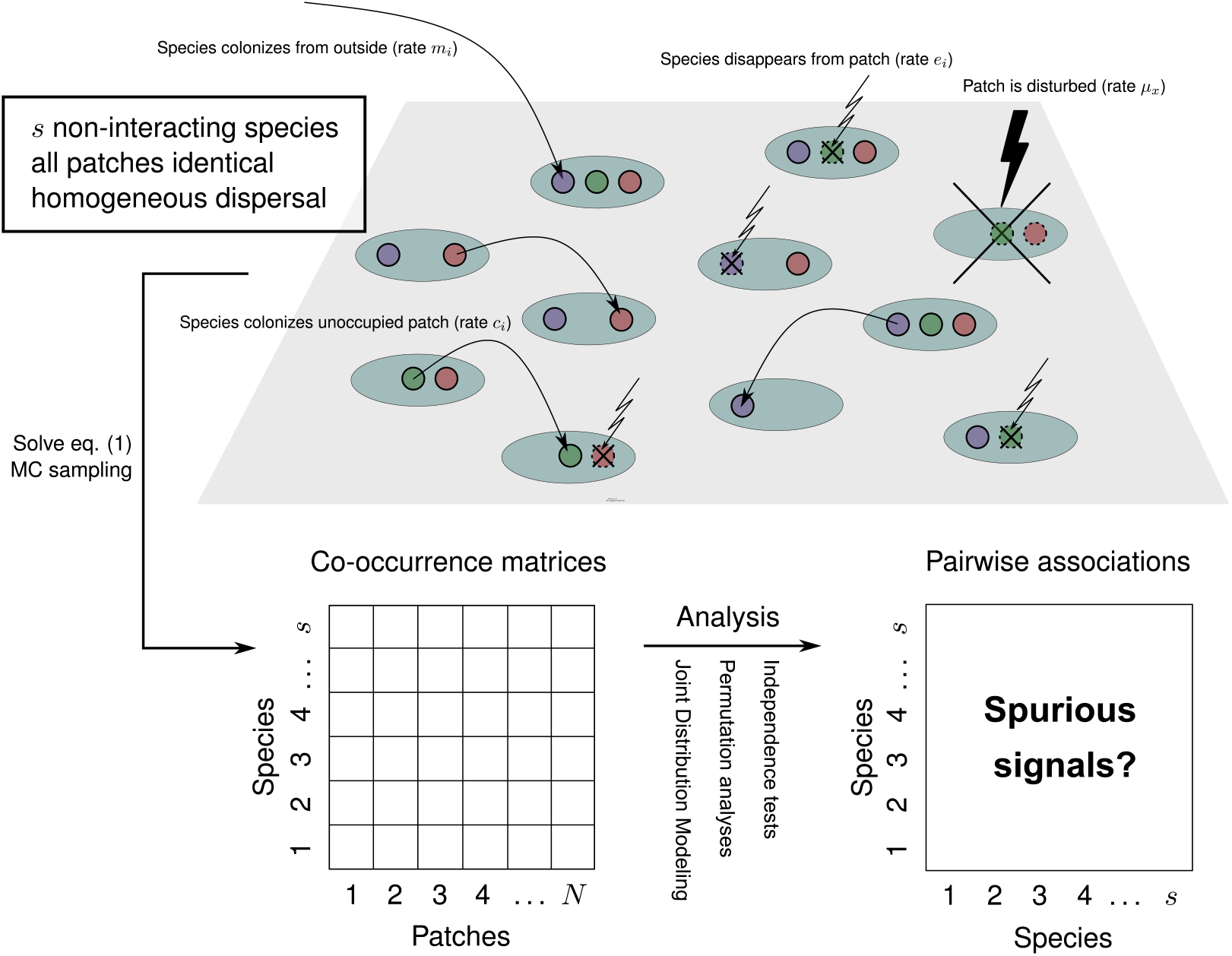
Graphical summary of our model and approach. Top: null metacommunity model, its assumptions and the processes it describes. Note that patch disturbances may occur at a certain rate, possibly dependent on the time since the last disturbance. Alternatively, they may occur after some prescribed amount of time (introducing a maximum life span of patches). Bottom: from the metacommunity model, using specific parameter values, we can generate sample co-occurrence matrices of arbitrary size. We can then subject these matrices to standard analytical methods to search for signals of species association. Considering that our model assumes no species interaction, no habitat heterogeneity or differences in habitat selection among species, we expect no significant association for any pair of species. Any such association would be a spurious signal (type I error, i.e. false positive).

Let *p_i,x,t_* be the fraction of patches that have age *x* and are occupied by species *i* at time *t* (Hastings, 1991). The fraction of patches occupied by species *i* is *p*_•,*x,t*_, and the fraction of patches with age *x* at time *t* is *p*_•,*x,t*_. The fraction of patches with age *x* that are not yet occupied by species *i* is therefore *p*_•,*x,t*_ – *p_i,x,t_*. We refer to Supporting Information A (section A.1.1) for more technical definitions. The general metacommunity model can be written as, for all *i* = 1, 2 *s*,

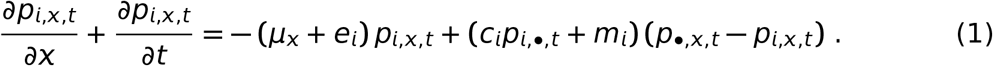

This framework generalizes classical metacommunity models (e.g. Cohen, 1970; Slatkin, 1974; Hastings, 1980; Hanski, 1982; Hastings, 1987; Tilman, 1994; Tilman et al., 1994; Hanski and Gyllenberg, 1997; Bell, 2001), deriving from MacArthur and Wilson (1963)’s “mainland-island” model (*c_i_* = 0), and from Levins (1969)’s metapopulation model (*m_i_* = 0). Such metacommunity models, when they consider patch catastrophic extinctions at all, assume a constant disturbance rate *μ_x_* = *μ*, but this need not be the case. The rate of disturbance might be increasing with patch age, producing an accelerating failure time model, for instance if a patch is a host that ages and suffers higher mortality with aging; this is termed a Type 1 survivorship curve in Begon and Townsend (2020). Conversely, if patches have some variability in their risk of disturbance, younger patches would go extinct at a relatively high rate, whereas older patches are relatively more disturbance resistant, causing a decline of *μ_x_*; with patch age (Type 3 survivorship curve). Another common situation is when patches have a finite lifetime and systematically get destroyed after some prescribed duration *X* (Pickett and White, 2013), as may happen for instance in agricultural settings where crops are harvested after some fixed time. Our framework encompasses all such situations. As special cases, the classical “mainland-island” and “Levins” models correspond to *c_i_* = 0 and *m_i_* = 0, respectively (Gotelli, 1991; Hanski and Gyllenberg, 1993), together with *X* → *∞* and *μ_x_* = *μ*, a constant (see Supporting Information A, section A2).

### 2.2 Steady-state occupancies and co-occurrence patterns

At steady-state, we can drop the *t* subscripts. The overall occupancy of species *i*, *p*_*i*,•_, can be derived explicitly in the special cases of the mainland-island or Levins models, for which simple expressions exist (Supporting Information A, section A2.1). For instance, *p*_*i*,•_ = 1 – (*e_i_* + *μ*)/*c_i_* in the Levins model. In general, no such solution exists when the rate of patch disturbance depends on age, but *p*_*i*,•_ can be computed numerically, e.g. by using a recursive algorithm outlined in Supporting Information A, section A1.4.

It would be possible to derive a separate equation for the dynamics of every possible patch state, i.e. the fraction of patches that are occupied by any given combination of species, and to solve these (as done in Supporting Information A, section A2.2, for two species). However, this is computationally inefficient. To obtain random co-occurrence matrices drawn from the above model, we instead use a Monte-Carlo procedure that generates the desired number of patches and draws their species composition. We provide a comprehensive set of R functions that make it easy to compute the steady-state occupancy of each species and generate random species co-occurrence matrices from the general model (1), for any number of species, patches, and any shape of the disturbance function *μ_x_* (Supporting Information B). We will use this method to generate null co-occurrence matrices that involve no species interactions, just metacommunity dynamics (Fig. 1).

### 2.3 Introducing relative distribution profiles and ‘fastness’

As illustrated in Fig. 2A, at steady state there exists a stable distribution of patch ages, and species occupancies vary as a function of patch age. We here derive a quantity *π*_*i*/*x*_ representing how species *i* is distributed over patch ages. We term this quantity the relative distribution profile of a species (Fig. 2B). Values smaller (larger) than one mean the species *i* is rarer (more frequent) in patches of age *x*, relative to its overall occupancy *p*_*i*,•_.

**Figure 2:**
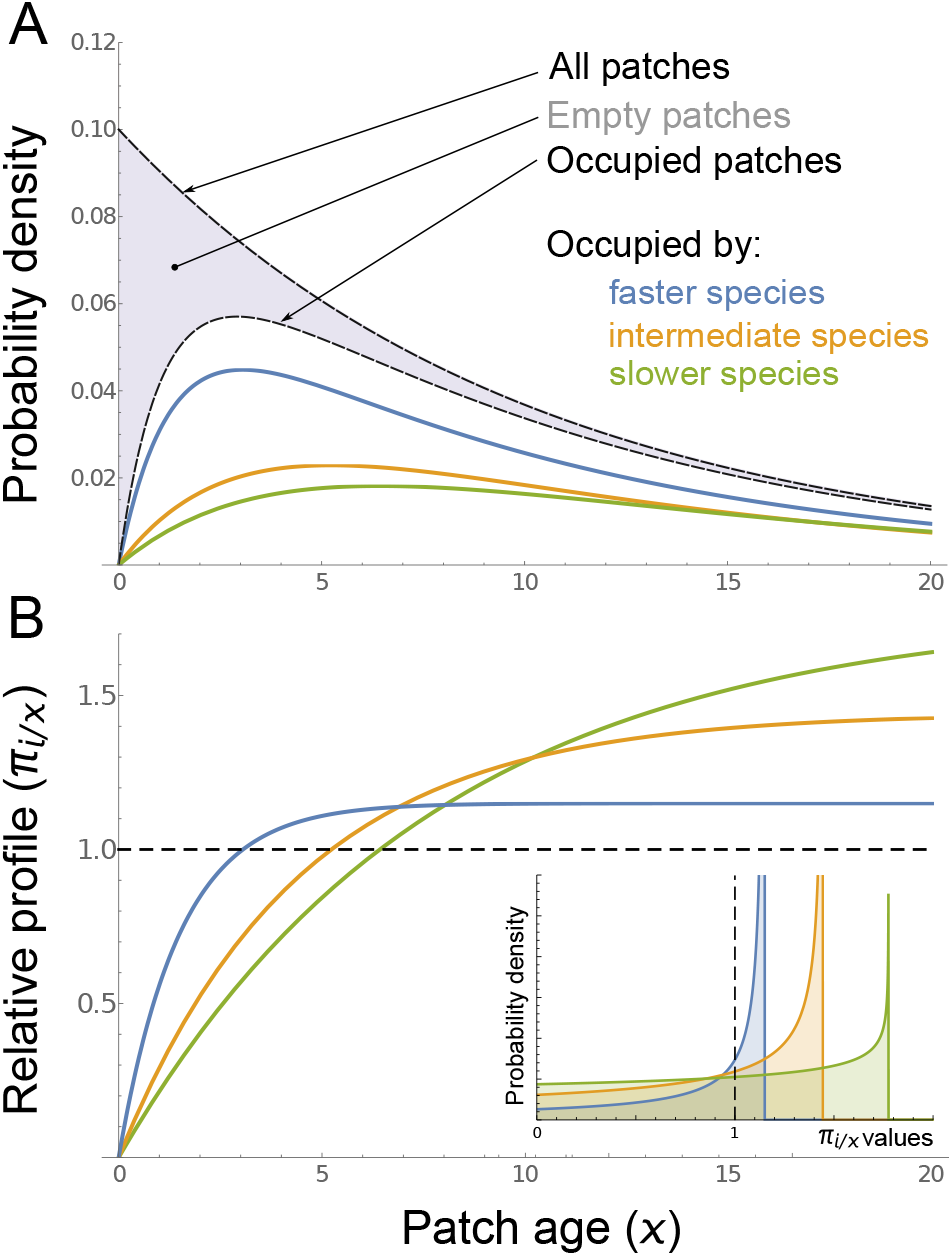
(a) Example equilibrium patch occupancies with three species differing in fastness. In this example the patch disturbance rate was *μ* = 0.1 per unit time with no maximum age, so that the stable patch-age distribution was exponential with mean 10 (top curve). At each patch age, all three species have some occupancy *p_i,x_* (bottom three curves), leaving a certain fraction of patches empty. In this example the fastest species are also more abundant overall, but this need not be the case. (b) The relative distribution profile of faster species has steeper initial slope and attains lower asymptotic values. As a species gets faster and faster, its profile converges to the horizontal dashed line, with value 1 for all patch ages. As shown in the insert, faster species have a smaller variance in *π*_*i*/*x*_ values, and as a species gets faster and faster, the distribution converges to a Dirac delta function at value 1 (vertical dashed line). Parameters: *m_i_* = 0.01, and *e_i_* = (0.05,0.1,0.2) and *e_i_* = (0.2, 0.3,1) for slow, intermediate and fast species, resp.

We first define the fraction of patches of age *x* that are occupied by species *i* as *p*_*i*|*x*_ = *p_i,x_*/*p*_•,*x*_, for which we obtain an explicit expression from model 1 at steady-state (Supporting Information A, section A1.3). The relative distribution profile of species *i* is then derived as the fraction of patches of age *x* that are occupied by species *i*, relative to its overall probability of occupancy *p*_*i*,•_:

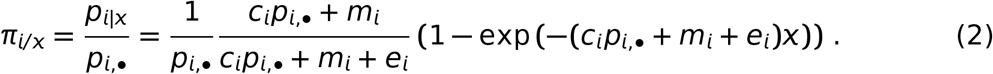

As shown in Fig. 2B, the mean value of the profile, for any species, is equal to one by definition (Supporting Information A, section A1.5).

To facilitate interpretation, we will discriminate species based on their “fastness”: fast species have very flat relative distribution profiles that quickly jump from zero and saturate at a value close to one (Fig. 2B). Slow species, on the contrary, have relative distribution profiles that slowly increase from zero and reach much larger values as the patch age gets large.

Formally, we will quantify the fastness of a relative distribution profile as its variance over patch ages, Vαr(*π_i_*/*x*). A maximally fast species has *π*_*i*/*x*_ = 1 for all *x*, i.e. zero variance (Fig. 2). Conversely, a very slow species has a relative distribution that gradually climbs to very large values, implying a huge variance. It can be shown that the variance of a species is also strictly decreasing with the initial slope of its relative distribution profile (Supporting Information A, section A1.7). The variance of a species profile is thus a metric of “slowness”, and its inverse is a metric of fastness. Importantly, the fastness of a species is not directly related to its overall occupancy in the metacommunity.

### 2.4 Methods to test for species associations in co-occurrence matrices

Methods to infer species associations from co-occurrence patterns are all based on the probability of co-occurrence, i.e. the fraction of patches in which both of the two species are found. Various methods exist to determine whether the observed probability deviates from statistical independence, under some appropriate “null” model. The probability of co-occurrence of two species *i* and *j*, i.e. the overall fraction of patches in which the two species are found, will be denoted *q*_*i,j*,•_ (at steady-state).

A straightforward null expectation is that of species independence: if species are not interacting, they should be distributed independently across patches, and thus the value of *q*_*i,j*,•_ should be compared to the null value *p*_*i*,•_*p*_*j*,•_ using a standard test for contingency tables (Veech, 2013). In this article we will use pairwise Fisher tests for independence as an exemplar of this type of method.

The prediction of species independence is straightforward and intuitive, but is known to suffer from several limitations. To circumvent these, more sophisticated null models are often preferred. A widespread approach consists of using permutation schemes intended to break species associations in co-occurrence matrices while retaining important species and patch differences (Gotelli, 2000). In this context, the C-score (Stone and Roberts, 1990) is the metric most commonly used to quantify the tendency of species to be segregated or aggregated in co-occurrence matrices. The partial C-score between two species, *C_i,j_* can be expressed as

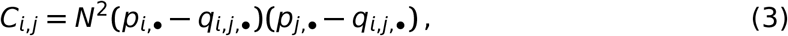

with *N* the total number of sites (patches) in the matrix.

Values of *C_i,j_* larger than expected indicate segregation (e.g. competition) between the two species, whereas values smaller than expected indicate preferential association (e.g. facilitative interactions). Several null models have been proposed to determine the expected *C_i,j_* value and test for deviations from it. The most classical methods are the so-called fixed-equiprobable (or sim2) and fixed-fixed (or sim9) permutation schemes, that reshuffle matrix values while keeping the row sums fixed, or both the row and column sums fixed, respectively (Gotelli and Ulrich, 2012; Münkemüller et al., 2020). We will consider both approaches.

More recently, flexible methods relying on the joint statistical modelling of species occurrences have gained a lot of popularity. These models are sufficiently flexible to allow several fixed or random predictors to be incorporated in modelling species occurrences, allowing different aspects of habitat variability or species trait differences to be controlled against. In such a framework, one can infer species associations from their residual covariances, i.e. the correlation between their residual probabilities of occurrence. One commonly used example of this approach are hierarchical models of species composition, as implemented in the Hmsc R package (Tikhonov et al., 2020; Ovaskainen and Abrego, 2020). This is the method we will use in this article.

In this framework, we will use four different model specifications. First, a null model (Hmsc M0) that includes no latent factor or covariate, and therefore is equivalent to the simplest methods above (Fisher’s tests or fixed-equiprobable permutation schemes). Second, differences across patches will be accounted for by including patch richness as a latent factor (Hmsc M1). Then, patch age (*x*) will be explicitly included as an environmental covariate in the model (Hmsc M2). As a variant of M2, patch richness will be also used as a proxy for patch age (Hmsc M2’).

### 2.5 Parameter values and numerical simulations

In the following, we will analyze how null co-occurrence matrices generated from our general metacommunity model behave, in terms of producing significant species associations, when subjected to the different methods described above (Fig. 1). We will first derive mathematical predictions, and test them with numerical simulations.

For all numerical simulations, we will use a synthetic community of 31 species, constructed to exhibit variation in their occupancies and extinction/colonization parameters (Fig. 4A; see Supporting Information B for details). Although the parameterisation does not correspond to a particular set of real species, the range of variation they present in colonization rates is similar to that found in natural metacommunities, for instance in Caribbean pond snails (Dubart et al., 2019) and grassland plant communities (Tilman, 1994).

**Figure 3:**
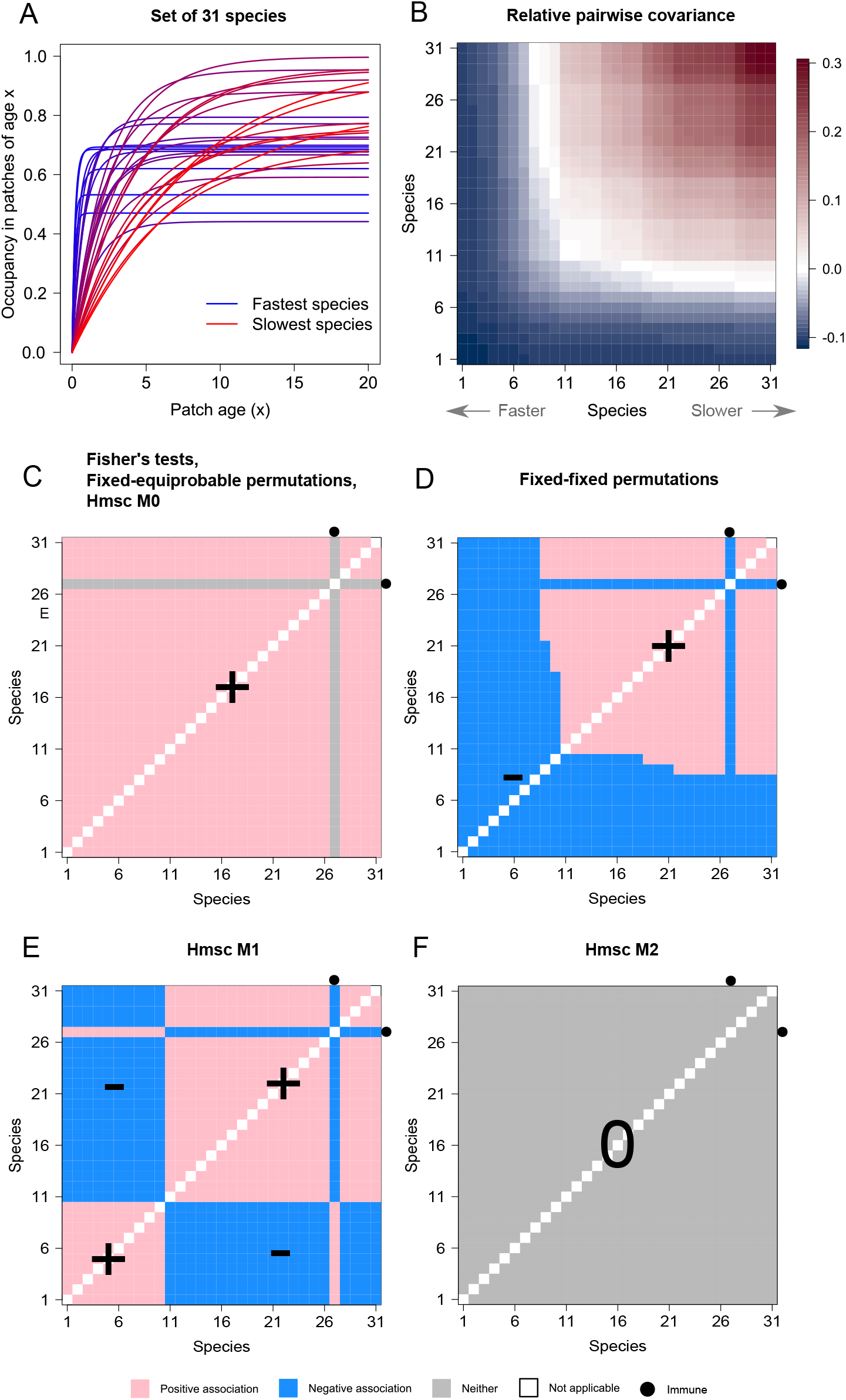
Mathematical predictions. (A) The set of synthetic species. Occupancy as a function of patch age *x* (*p_i|x_*) is shown for each of the 31 species. Species traits were drawn randomly in order to have variable overall occupancies and fastnesses. (B) Pairwise covariances of relative profiles for all species pairs. Species are ranked according to fastness: species 1 is the fastest (lowest variance), and species 31 is the slowest (highest variance). The variance of the average profile in the community 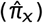 was substracted from covariances to help distinguish lower/higher than average values (color bar). See equations (4) and (7). (C) Mathematical predictions associated with methods testing for pairwise species independence (Fisher’s test, fixed-equiprobable-Sim2-permutations, and the simplest joint species distribution model Hmsc M0). (D–E) Same as (C) based on methods that control for patch richness, i.e. the fixed-fixed permutation (Sim9) algorithm (D), and joint species distribution model Hmsc M1 (E). (F) Same as (C) based on joint species distribution model Hmsc M2, a method that controls for patch age. Species are ordered as in (B). The bullets indicate the predictions relative to an example species (species 27) being immune to patch disturbance.

**Figure 4:**
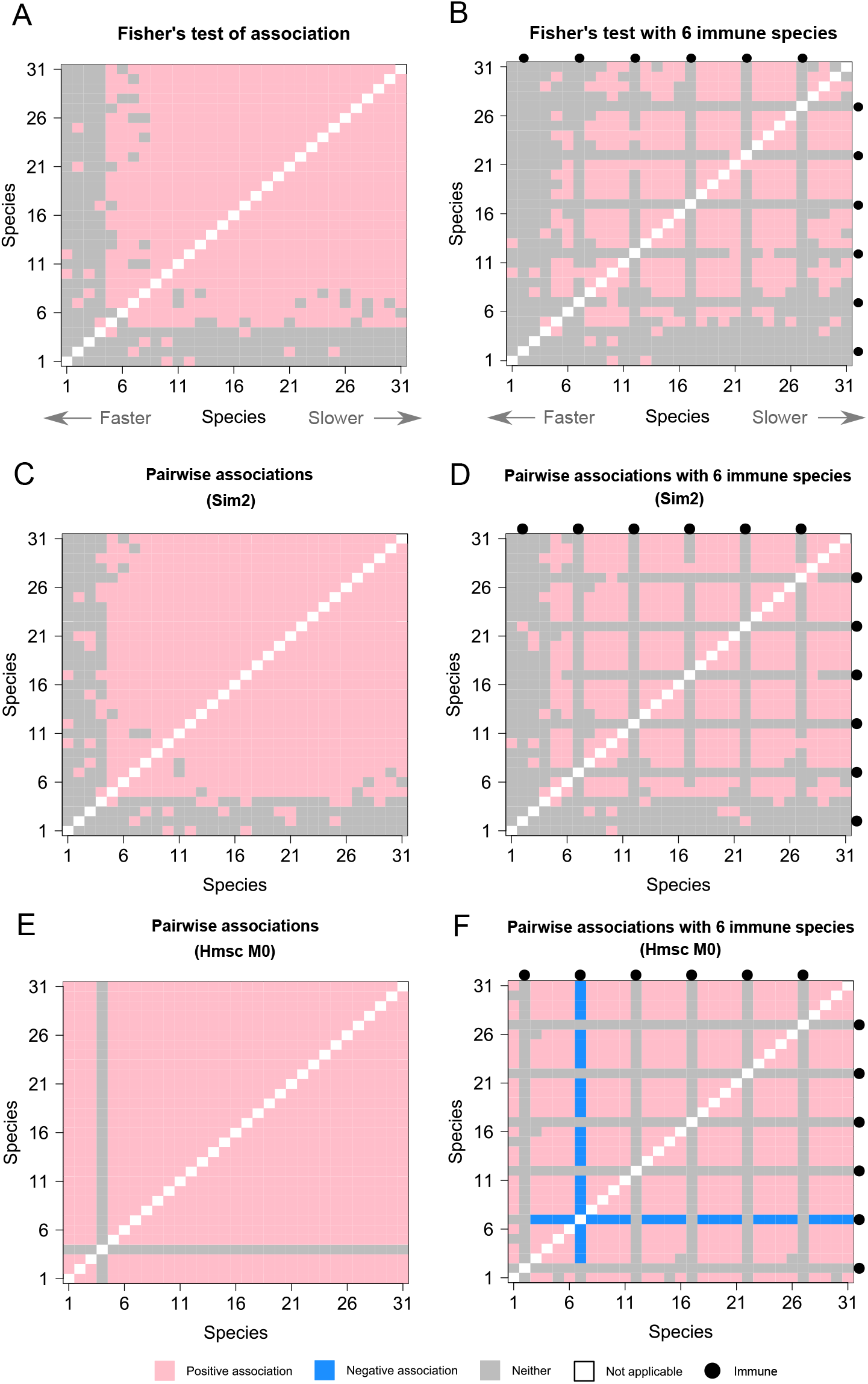
Numerical simulations: methods testing for pairwise species independence. (A) Species pairs significantly associated according to Fisher’s pairwise tests. (B) Same as (A), with a subset of species immune to patch disturbance (shown as bullets). (C) Species pairs significantly associated according to the fixed-equiprobable (Sim2) permutation algorithm. (D) Same as (C), with a subset of immune species. (E) Species pairs significantly associated according to the simplest joint species distribution model (Hmsc M0). (F) Same as (E), with a subset of immune species. Species are ordered as in Fig. 3B. The sample co-occurrence matrix was generated from equation (1) with *N* = 1000 sites (patches) and the illustrative parameter set.

We assumed the most classical metacommunity set-up, i.e. no external immigration (*m_i_* = 0) and a constant rate of patch disturbance. The value of the disturbance rate was set to *μ*(*x*) = 0.1 or *μ*(*x*) = 0.2 per unit of time. The maximum life-span of patches was set to *X* = 20 time units. These values correspond to low or intermediate levels of patch disturbance, considering the average species-specific extinction rate was 0.5 and the average colonization rate was 1.5. This means that extinctions caused by patch disturbance were always several times less frequent than other extinction/recolonization events. We chose these values in a conservative mindset, knowing that in metacommunities more dominated by patch disturbance, and/or with more variable species traits, the patterns we report could be more pronounced.

We varied the sample size (number of patches in the co-occurrence matrices) between 300 and 1500, with intermediate values 500 and 1000 (for recent studies using similar sample sizes, see Dubart et al., 2019; Opedal et al., 2020a; Facon et al., 2021). Finally, we also considered the possibility that not all species might be affected by patch disturbance: some species might be immune to disturbance (“immune species”). To address this possibility, we used the same set of 31 species, but rendered six of them, evenly spaced along trait values, invulnerable to patch disturbance (*μ*(*x*) = 0 for them), for each parameter combination.

This corresponded to a total of 2 × 4 × 2 = 16 parameter combinations. For each combination, we generated random co-occurrence matrices from the null metacommunity model (eq. 1). Each time, we applied the different methods introduced above to test for significant species associations, with standard settings recommended in the literature. We computed, for each parameter combination and statistical method, the percent of spurious positive associations (fraction of species pairs declared positively associated) and the percent of spurious negative associations (fraction of species pairs declared negatively associated).

All simulations and figures reported can be reproduced using the accompanying R markdown (Supporting Information B).

## 3 Results

### 3.1 Mathematical predictions

#### 3.1.1 Patch disturbance generates spurious positive associations with methods that do not account for patch age

##### Direct independence tests (e.g. Fisher’s tests)

In the absence of any interaction, species are independently distributed among patches within any particular age class (Supporting Information A, section A3). This implies

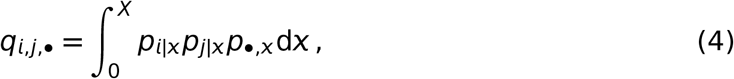

which can be equivalently expressed as:

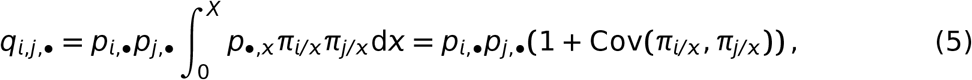

given that the mean of the profile is one (Supporting Information A, section A1.5). Since *π_i/x_* and *π_j/x_* are increasing functions of *x* (Eq. 2), from Harris’ inequality we have Cov(*π_i/x_*, *π_j/x_*) ≥ (Supporting Information A, section A1.5). From (5) we therefore have

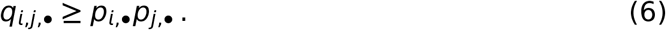

Equality occurs only if at least one of the two species is distributed uniformly over all patch ages, i.e. *π_k/x_* = 1 for all *x*, so that *Cov*(*π_i/x_*, *π_j/x_*) = 0. From Eq. 2, this is not the case in general. This only occurs if there is no patch disturbance at all (*μ_x_* = 0 and *X* → ∞), in which case all patches are effectively infinitely old, so that *π_i/x_* and *π_j/x_* both tend to one, and *Cov*(*π_i/x_*, *π_j/x_*) tends to zero. But in all other cases, species are not distributed independently: they are positively associated, i.e. they co-occur more often than expected from the “null” product of their respective overall occupancies (see Eq. 6).

As a consequence, methods based on the test of pairwise independence of species, such as direct Fisher’s tests or more recently proposed methods (e.g. Veech, 2013) would consistently detect spurious positive associations (aggregation) among species, even in the absence of any interaction between species. Specifically, positive associations will be strongest, and thus more likely to be statistically detected, for species pairs with a large covariance of their relative profiles (Eq. 5), i.e. for pairs of species that are both slow. In contrast, species pairs including at least one fast species are more likely to be declared independent.

##### Fixed-equiprobable permutations (Sim2)

The same conclusion holds for more elaborate methods that do not control for patch age, such as the “fixed equiprobable” permutation algorithm, also known as “Sim2” (Gotelli, 2000). To show this, let us consider the commonly-used partial C-score *C_i,j_* as our metric for species association (Eq. 3). We will denote with an asterisk the value of quantities after the permutation algorithm has been applied to a co-occurrence matrix.

A fixed-equiprobable matrix permutation algorithm preserves the number of occurrences per row (i.e. overall occupancies of species) but reshuffles species across all patches equiprobably. Therefore, after permutations, species overall occupancies are unchanged (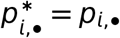 for all *i*), but all species become equiprobable over the different age classes, on average. Therefore, their relative distribution profiles become flat: 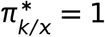 for all *x*. From Eq. 5 it follows

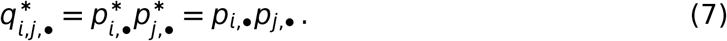

This indicates that the algorithm effectively breaks the positive association generated by patch disturbance. From Eq. 6, this implies 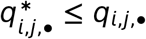. Permuted matrices will on average have a deficit of co-occurrences, for all species pairs, compared to the original matrix. Since *p*_*i*,•_ and *p*_*j*,•_ are unaffected by permutation, we can directly conclude that 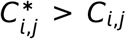, as soon as patch disturbances occur. Using the fixed-equiprobable permutation algorithm will therefore consistently yield spurious facilitation signals between virtually all pairs of species, exactly as direct tests for species independence do (see Fig. C1A in Supporting Information C).

##### Joint species distribution modelling (Hmsc M0)

Joint species distribution models that do not account for patch differences (such as our Hmsc M0), by assuming species to be independently distributed across patches, will suffer from the same caveat.

Hmsc models predict the probability of presence of some species *i* in a patch of age *x*, 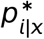, where the asterisk stands for “predicted value”.

The model predicts the probability that species *i* is present in a patch regardless of the patch age, i.e.

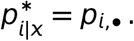

It follows that the residual association (covariance) of two species *i*, *j* is

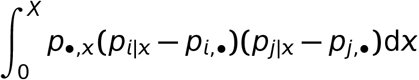

which is equal to, dividing by *p*_*i*,•_,*p*_*j*,•_,

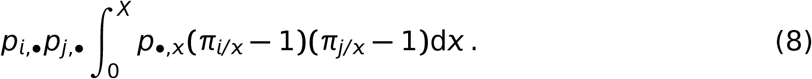

Since all species have relative distribution profiles smaller than one for small *x*, and greater than one for large *x*, it follows that the integrand is positive for most *x* values, and thus the two species will have positive residual association.

Note that if at least one of the two species is infinitely fast, then the corresponding parenthesis is null for all *x*: the two species will have no residual association (i.e. will be independently distributed).

Hmsc M0 models will thus report positive residual associations for all species pairs. The ability to detect those associations will however depend on statistical power, especially for pairs of fast species, as in previous methods (independence tests and fixed-equiprobable permutations).

#### 3.1.2 Patch disturbance yields a characteristic signature of spurious associations even with methods that control for patch richness

We now turn to methods that can control for systematic differences among patches, in this case emerging from heterogeneity in the time since the last disturbance occurred (patch age). We here distinguish two groups of approaches. We first treat the popular fixed-fixed permutation algorithm, also known as Sim9 or swap (Gotelli, 2000). We then consider joint species distribution models that include patch richness as a factor (as our Hmsc M1).

##### Fixed-fixed permutation approaches (Sim9)

By fixing the row totals, this method effectively maintains the number of species per patch unchanged, and can thus deal with the positive association of species that causes issues with the previous set of approaches.

Similar to the permutation algorithm discussed in the previous section (Sim2), preservation of row totals implies 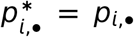. Therefore, we only need to study changes in *q*_*i,j*,•_ in order to understand changes in C-scores. Preservation of column totals further means that the number of occurrences will remain the same in every age class, which can be stated as 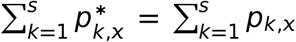 for all *x*. Species are otherwise reshuffled indifferently across all patch ages. Let *w_i_* be the relative occupancy of species *i* in the matrix, that remains unchanged after permutations:

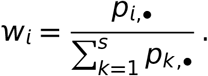

A fixed-fixed permutation algorithm yields:

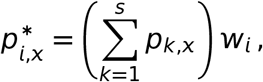

leading to, for all *i* = 1, 2… *s*,

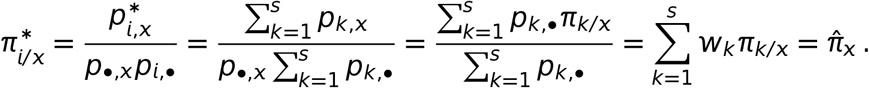

This indicates that species relative distribution profiles are replaced with their weighted-average over all species, with weights equal to species relative overall occupancies. Let us denote this weighted-average by 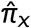. We note that 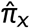 can also be expressed as

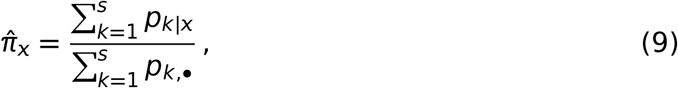

i.e. 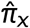 is the mean species richness in patches age of *x* relative to the mean richness over all patch ages.

We thus obtain:

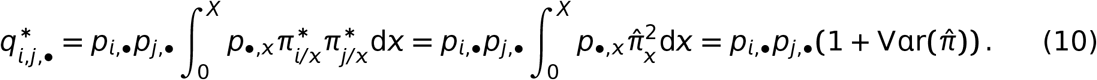

Comparing with Eq. 5, we see that the term correcting for non-independence, Cov(*π_i/x_*, *π_j/x_*), is here replaced with 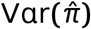. The difference 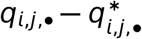 has the same sign as 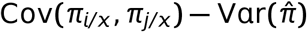.

The permutation algorithm is thus biased, except if 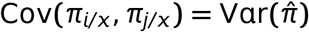 for all species pairs. This would occur if all species had identical relative distribution profiles, i.e. when all species are “similar”, in the sense *p_i,x_* = *κp_j,x_* for all *x*, with some positive *κ*. This trivially occurs if there is no patch disturbance at all, or if all species have identical extinction and colonization parameters. Beyond that, species may differ in their parameters and overall occupancies and still be similar in that sense, but this require quite constrained parametric conditions (Supporting Information A, section A2.3).

Except for these restricted scenarios, the permutation algorithm will yield spurious associations in both directions. Species pairs whose relative distribution profiles have covariance smaller than the variance of the average community profile 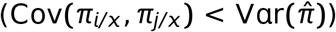 will appear to have a spurious negative association. Conversely, species pairs for which 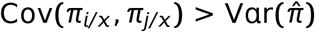 will appear to have a spurious positive association. Only species pairs for which 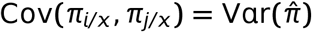 would appear as independently distributed.

In practical terms, species pairs containing at least one species that is ‘faster’ than average will have low relative covariance, and thus will be declared negatively associated. Indeed if the fast species (say *i*) is fast enough (regardless of how slow the other is), then *π*_1/*x*_ ≈ 1 for all *x* thus and Cov(*π_i/x_*, *π_j/x_*) ≈ 0 = Vαr(*π_i/x_*), which is less than the variance of the average. In contrast, species pairs that are both ‘slower’ than average will have large relative covariance, and will be declared positively associated. Pairs of species that are both close to average fastness may yield no particular signal.

If species are ordered according to fastness (i.e. variance), this produces a characteristic signature of spurious association signals. The pairwise covariance of all species pairs used in reference set of species is shown in Fig. 3B. The corresponding mathematical predictions for fixed-fixed permutation algorithms are illustrated in Fig. 3D (note the vertical and horizontal blue lines for species 27 are because that species is assumed to be immune to patch disturbance as an illustration of the potential effect of that phenomenon; see below).

##### Joint species distribution modelling (Hmsc M1)

In the context of joint species distribution modelling, it has been suggested that an equivalent to the fixed-fixed permutation scheme discussed above would be to include patch richness as a latent factor in the model of species occurrence (our Hmsc M1). This was hypothesized in (Ovaskainen and Abrego, 2020, p. 335; see Supporting Information B for details).

We can similarly formulate mathematical predictions regarding the behavior such an approach would have under our null metacommunity model. In the limit of large co-occurrence matrices, the number of species in a patch (Σ*_k_p_k|x_*) tends to become tightly associated with patch age *x*. In other words, one may neglect the variance in patch age for a given species richness. A statistical model that describes the probability of occurrence, while including patch richness as a latent factor, would therefore predict species *i* to occur in some particular patch of age *x* with probability 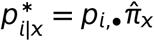, since 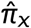 is the relative richness of a patch of age *x* (Eq. 9).

The expected residual for this species in this sort of patch would be 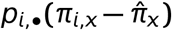, and the residual covariance between two species *i* and *j*, over all patches, would be, from Eq. (8),

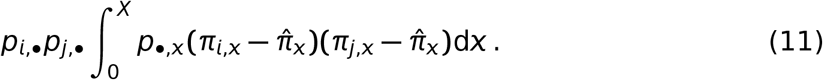

From the above equation we can immediately conclude that if at least one of the two species has average fastness, in the sense defined above, then 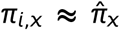 for all *x*, yielding a residual association of zero (independence). Otherwise, if the two species are faster (slower) than average, then both parentheses in the integral will be first positive (negative), and then negative (positive), as *x* increases. It follows that the overall residual association will be found positive. By the same argument, if one species is faster than average and the other is slower, the two parentheses will have opposite signs, and the overall residual association will be negative.

As for the above permutation scheme, we expect a mixture of positive and negative associations, unless all species have the same fastness. The average fastness again plays a critical role in determining where the switch occurs. However, the resulting pattern of species associations is markedly different. We predict a block-like pattern, with faster species positively associated together, slower species also positively associated, and the two groups of species mutually negatively associated. We thus predict a four-block pattern, reminiscent of differential specialization on two habitats, as illustrated in Fig. 3E.

In any case, controlling for differences across patches is therefore insufficient to handle the metacommunity dynamical consequences of patch disturbance, even though it brings some improvement. First, correct results are obtained if all species are “similar” (i.e. have identical fastnesses). Second, the overall number of spurious associations obtained will likely be reduced, since some species pairs will likely be correctly declared as independently distributed. However, this comes at the price of generating spurious associations of both kinds (positive and negative), not only positive associations.

#### 3.1.3 Patch disturbance requires species-specific modelling of patch-age

Efficiently dealing with patch disturbance in co-occurrence analyses therefore requires both patch age and species traits to be taken into account, in order to model appropriately the differential effect that patch-age has on species with different fastnesses. In other words, there must be an interaction between the effect of patch-age and the identity of species.

This cannot be done with direct independent tests or simple null model approaches, but can be in the context of joint species distribution models, such as our model Hmsc M2.

The model predicts the probability that species *i* is present in a patch as an increasing function of patch age, i.e. it will yield 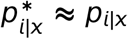 where the approximation denotes the fact that the functional form of the regression curve may not exactly match the shape of the *p_i/x_* function.

It follows that residuals will be close to zero, and so will be the residual associations, for all species pairs. This is illustrated in Fig. 3F.

#### 3.1.4 What if some species are immune to patch disturbance?

So far, we assumed all species in the metacommunity were affected by patch disturbances. In practice, in large sets of species, it may be that only a subset of species are susceptible to disturbance events, while some others are not (“immune” species). Based on our mathematical results, this possibility is straightforward to deal with: the immune species would behave as a species with infinite fastness (Vαr(*π_i/x_*) = 0), regardless of its actual colonization/extinction rates (Supporting Information A, section A4).

As a consequence, any species pair including at least one immune species is expected to return no overall association, if using pairwise independence tests or equivalent methods treated in Section 2.1; see Fig. 3C). Therefore, fewer spurious associations would be detected with such methods.

On the contrary, by increasing the heterogeneity of fastness among species in the community, the presence of immune species would degrade the performance of methods that control for patch richness. More specifically, when using a fixed-fixed permutation method, any species pair including one immune species will systematically appear as negatively associated with the others, since Cov(*π_i/x_*, *π_j/x_*) = 0. For the same reason, if using a joint species distribution model with patch richness as a latent factor, immune species will tend to cluster with the fastest species, and to be declared negatively associated with the slower species (Fig. 3D–E).

All the mathematical predictions derived in this section are summarized in Fig. 3. We will now proceed to test them with numerical simulations.

### 3.2 Numerical simulations

#### 3.2.1 Patch disturbance generates spurious positive associations with methods that do not account for patch age

Consistent with our mathematical predictions, species were positively associated overall in the simulated co-occurrence matrices, even under our “null” model (eq. 1). Using the reference parameter set (*μ* = 0.1, *N* = 1000 and no immune species), the odds of species co-occurrence were on average 75% higher than expected under independence (Table 1). About 10% of all species pairs had odds of co-occurrence more than 2.5 times as large as expected.

**Table 1:** Performance (% of errors) for each of the 5 methods tested (Fisher, Fixed-fixed, Hmsc M0, Hmsc M1, Hmsc M2), under the 16 parametric scenarios (sample size *N* x disturbance rate μ x No Immune species/With immune species). Note that the “fixed-fixed” algorithms is also known as “Sim9”.

| Scenario | Fisher |  |  | Fixed-fixed |  |  | Hmsc M0 |  |  | Hmsc M1 |  |  | Hmsc M2 |  |  |
| --- | --- | --- | --- | --- | --- | --- | --- | --- | --- | --- | --- | --- | --- | --- | --- |
|  | %+ | %- | %tot | %+ | %- | %tot | %+ | %- | %tot | %+ | %- | %tot | %+ | %- | %tot |
| **No immune species** |  |  |  |  |  |  |  |  |  |  |  |  |  |  |  |
| $\mu = 0.2$ $N = 300$ | 38.3 | 0 | 38.3 | 11.8 | 8.8 | 20.6 | 75 | 0 | 75 | 0 | 0 | 0 | 0 | 0 | 0 |
| $N = 500$ | 58.3 | 0 | 58.3 | 20.2 | 19.8 | 40 | 78.3 | 0 | 78.3 | 0.6 | 3.9 | 4.5 | 0 | 0.4 | 0.4 |
| **\* $N = 1000$** | **73.8** | **0** | **73.8** | **29.7** | **27.3** | **57** | **98.9** | **0** | **98.9** | **4.1** | **8** | **12.1** | **0.9** | **2.8** | **3.7** |
| $N = 1500$ | 79.6 | 0 | 79.6 | 35 | 28.6 | 63.6 | 98.5 | 0 | 98.5 | 6.9 | 13.8 | 20.6 | 1.3 | 1.9 | 3.4 |
| $\mu = 0.1$ $N = 300$ | 19.6 | 0 | 19.6 | 5 | 3.6 | 8.6 | 69.9 | 0 | 69.9 | 0 | 0.2 | 0.2 | 0 | 0 | 0 |
| $N = 500$ | 45.2 | 0 | 45.2 | 11.8 | 10.5 | 22.4 | 70.1 | 0 | 70.1 | 0.4 | 1.1 | 1.5 | 0 | 0.6 | 0.6 |
| $N = 1000$ | 69.4 | 0 | 69.4 | 19.1 | 18.5 | 37.6 | 98.5 | 0 | 98.5 | 0.9 | 2.5 | 3.4 | 0 | 0 | 0 |
| $N = 1500$ | 75 | 0 | 75 | 25.6 | 22.6 | 48.2 | 87.3 | 0 | 87.3 | 2 | 6.4 | 8.4 | 0 | 0.2 | 0.2 |
| **With immune species** |  |  |  |  |  |  |  |  |  |  |  |  |  |  |  |
| $\mu = 0.2$ $N = 300$ | 22.6 | 0 | 22.6 | 4.7 | 6 | 10.7 | 50 | 0 | 50 | 0 | 0 | 0 | 0 | 0 | 0 |
| $N = 500$ | 37.8 | 0 | 37.8 | 20.4 | 17.8 | 38.3 | 45.2 | 0 | 45.2 | 1.1 | 7.3 | 8.4 | 0 | 0 | 0 |
| $N = 1000$ | 48.4 | 0 | 48.4 | 35.3 | 23.4 | 58.7 | 64.1 | 4.7 | 68.8 | 9.9 | 18.5 | 28.4 | 0 | 0.4 | 0.4 |
| $N = 1500$ | 51.8 | 0 | 51.8 | 41.3 | 27 | 68.4 | 59.3 | 0 | 59.3 | 5.6 | 20 | 25.6 | 0 | 0 | 0 |
| $\mu = 0.1$ $N = 300$ | 7.5 | 0 | 7.5 | 1.9 | 3.6 | 5.6 | 45.2 | 0 | 45.2 | 0 | 0 | 0 | 0 | 0 | 0 |
| $N = 500$ | 28.2 | 0 | 28.2 | 10.1 | 10.3 | 20.4 | 45.2 | 0 | 45.2 | 3 | 5.8 | 8.8 | 0 | 0 | 0 |
| $N = 1000$ | 44.7 | 0 | 44.7 | 20.4 | 19.8 | 40.2 | 64.5 | 3.9 | 68.4 | 3.2 | 7.3 | 10.5 | 0 | 0 | 0 |
| $N = 1500$ | 49.2 | 0 | 49.2 | 27.7 | 24.9 | 52.7 | 64.5 | 3.9 | 68.4 | 3.2 | 7.3 | 10.5 | 0 | 0 | 0 |
| **Grand average** | 46.8 | 0 | 46.8 | 20 | 17 | 37 | 69.6 | 0.78 | 70.4 | 2.5 | 6.3 | 8.9 | 0.17 | 0.23 | 0.33 |

\* This is the scenario used in all figures.

Results for methods not controlling for patch richness or patch age (pairwise Fisher’s tests, fixed-equiprobable permutations, and species distribution model Hmsc M0) are shown in Fig. 4. As predicted, all methods detected significant positive associations for the vast majority of species pairs, except for some of the species pairs involving the fastest species. Direct pairwise Fisher’s tests and fixed-fixed permutations yielded very similar results, with about 75% of species pairs declared as positively associated (Fig. 4A,C). Hmsc M0, that uses a quite different statistical approach, happened to be more sensitive, and thus more error prone, declaring more than 90% of species pairs positively associated (Fig. 4E).

Again confirming predictions, the inclusion of species immune to patch disturbance reduced the number of positive associations detected, as species pairs including at least one immune species were reported as independently distributed, with a few exceptions (Fig. 4B,D,F). The main difference between the three methods was that Hmsc M0 could occasionally go as far as declaring one immune species as negatively associated with the non-immune species (Fig. 4F).

#### 3.2.2 Patch disturbance yields a characteristic signature of spurious associations even with methods that control for patch richness

Simulation results applying fixed-fixed permutation schemes (Sim9) and the species distribution model Hmsc M1 are presented in Fig. 5. As expected from our mathematical analyses, we observe the detection of spurious associations, of both kinds (positive and negative). The patterns of species associations closely resemble those of mathematical predictions, be it for fixed-fixed permutations (see Fig. 3D) or Hmsc M0 (see Fig. 3E). The main difference from predictions is the absence of some predicted associations, due to lack of statistical power. This is particularly the case for species pairs with covariance close to average, for a fixed-fixed permutation algorithm (Fig. 5A).

**Figure 5:**
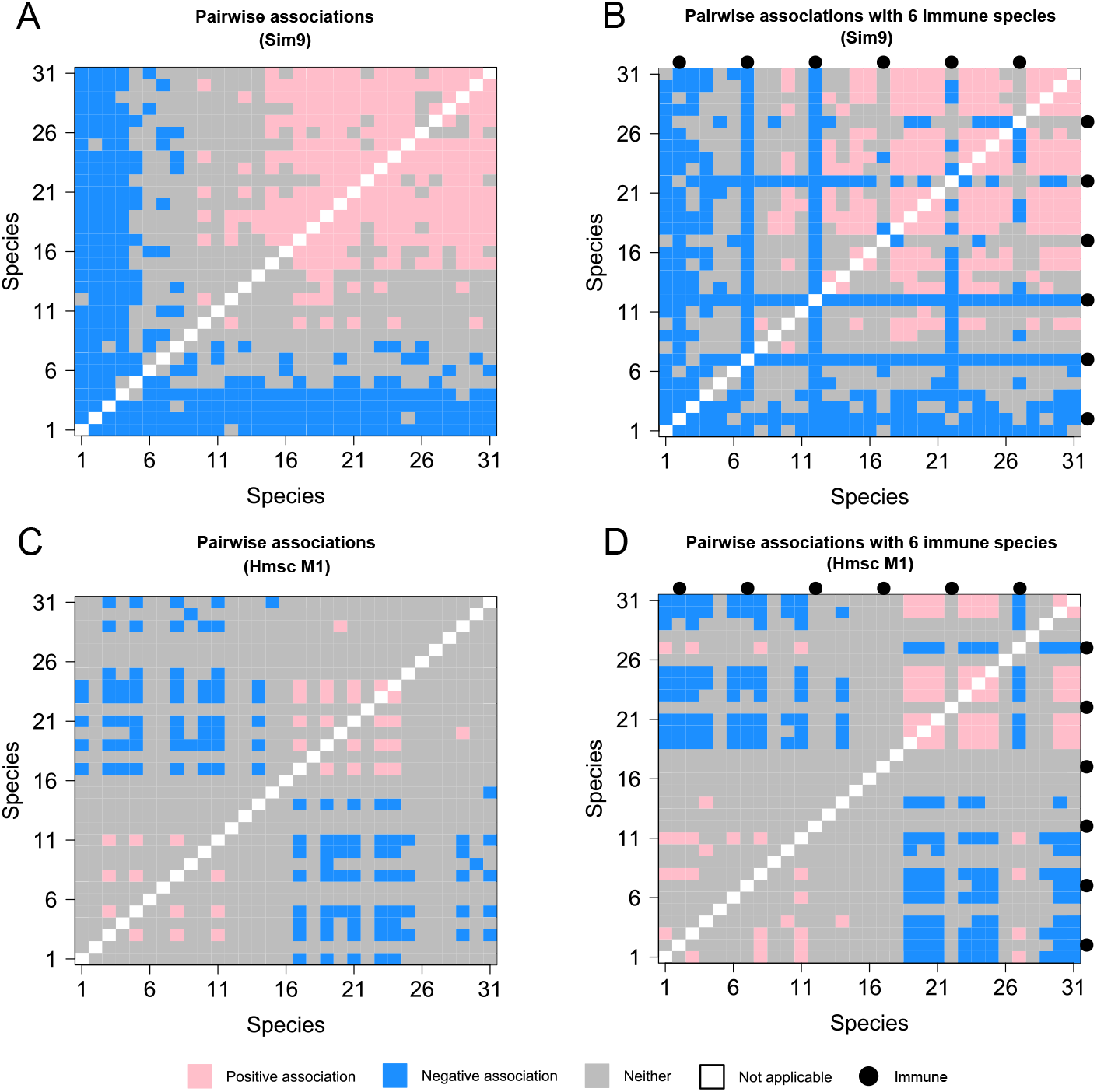
Numerical simulations: methods controlling for patch richness. (A) Species pairs significantly associated according to the fixed-fixed (Sim9) permutation algorithm. (B) Same as (A), with a subset of species immune to patch disturbance (shown as bullets). (C) Species pairs significantly associated according to a joint species distribution model (Hmsc M1) that models patch richness as a single latent factor. (D) Same as (C), with a subset of immune species. Same species and sample co-occurrence matrix as in previous figures.

The total number of species pairs for which a spurious association is reported is slightly smaller than it was without controlling for patch richness. It is 56% with a fixed-fixed permutation scheme versus 73% with a fixed-equiprobable scheme. Similarly, it is 12% with Hmsc M1 versus more than 90% with Hmsc M0.

However, both positive and negative spurious associations are detected. There is a slight majority of negative associations with fixed-fixed permutations, but, in contrast, a majority of positive associations with Hmsc M1. Unlike what was observed for the previous set of methods, Hmsc modelling is here less sensitive, and thus less error prone, than the comparable permutation algorithm. Therefore, fixed-fixed permutations and Hmsc M1 models differ not only in the qualitative type of characteristic signature they produce, but also in their quantitative performance (power and ratio of negative versus positive associations).

Introducing immune species also yielded results conforming to mathematical predictions (Fig. 5B,D). Interestingly, the presence of immune species degraded the overall performance of the two methods, but not to the same extent: the percentage of spurious associations detected increased from 12% to 28% in the case of Hmsc M1 and, more modestly, from 57% to 59% in the case of fixed-fixed permutations.

#### 3.2.3 Patch disturbance requires species-specific modelling of patch-age

Species distribution models can describe how the probability of occurrence increases, for each species, as a function of patch age. This can be done rather straightforwardly by providing patch-age as an environmental covariate, over which species occupancies are regressed, as in our model Hmsc M2.

As predicted, such an approach can restore satisfying results, i.e. no association was detected. Examples are shown in Fig. 6A–B. Using the same set of species and co-occurrence matrix as before, the use of model Hmsc M2 suppressed almost all spurious association signals, with or without immune species, as desired. The percentage of spurious associations detected was not strictly speaking zero, but remained under a tolerable error rate of 5%.

**Figure 6:**
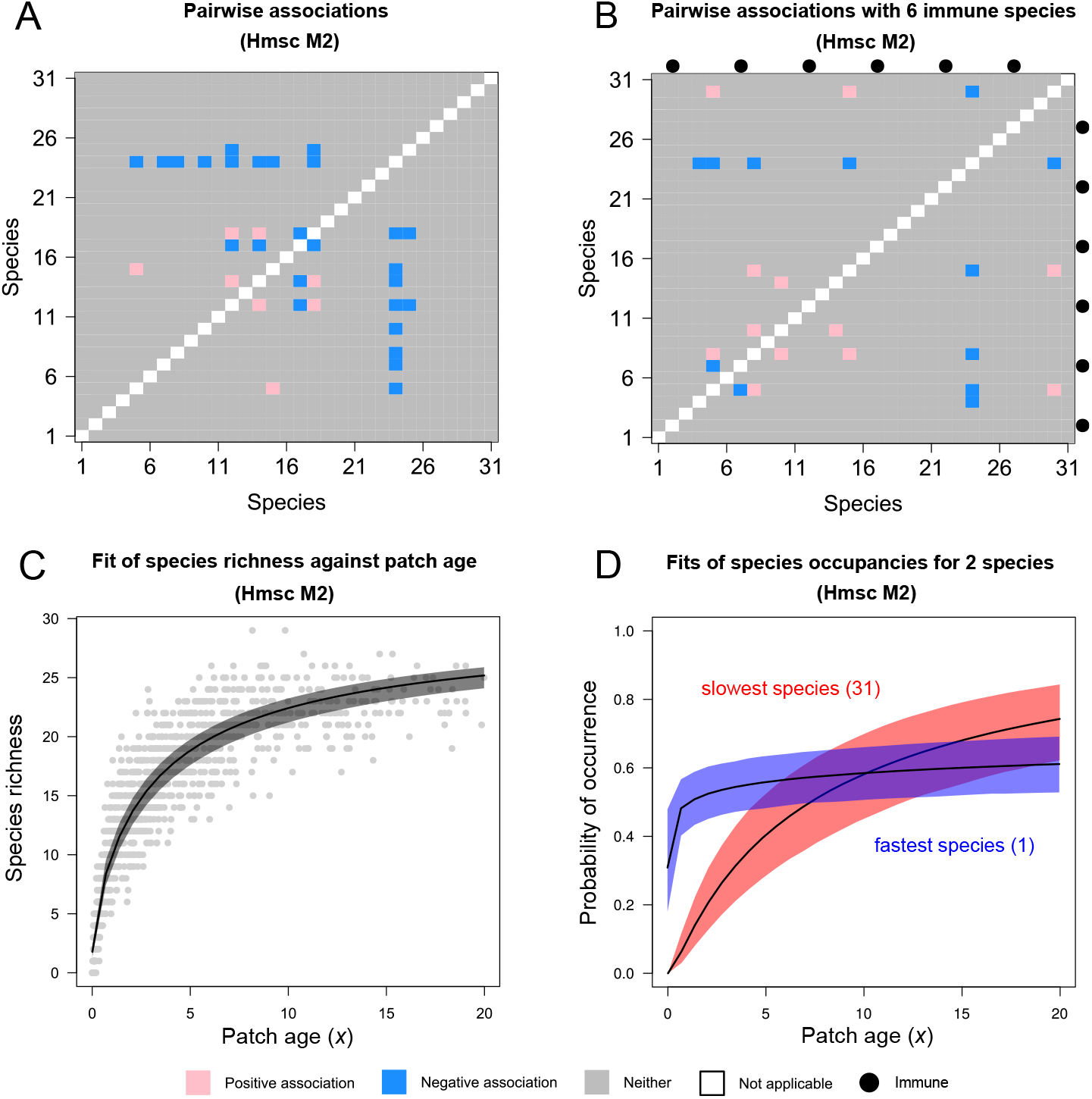
Numerical simulations: modelling the effect of patch age. (A) A joint species distribution model that incorporates patch-age as an explicit environmental covariate (Hmsc M2) yields correct results, even with patch disturbance. The few spurious species associations detected are within the type-I error rate. Note that log(patch age) was used as the covariate, since patch age is distributed exponentially. (B) Same as (A), with a subset of species immune to patch disturbance (shown as bullets). (C) Predicted species richness as a function of patch-age. Gray dots represent the actual data in the sample co-occurrence matrix. (D) Predicted probability of occurrence as a function of patch age, for the fastest species and the slowest species. Same species and sample co-occurrence matrix as in previous figures.

By regressing the probability of occurrence on patch age, for every species individually, such a model appropriately captures both the overall increase in occupancy with patch age (Fig. 6C) and the differential response of the different species, depending on their fastness (Fig. 6D).

Note however that this approach requires additional data, namely the age of patches, on top of the species co-occurrence matrix. This would not always be possible. Considering the fair association between patch age and patch richness (Fig. 6C), one could consider using patch richness as a surrogate for patch age. As patch richness is already encoded in the species co-occurrence matrix, this approach, like all previous sets of methods, would not require additional data.

However, if using patch richness as a proxy of patch age (model Hmsc M2’), the method fails to provide correct results (Supp. Fig. S1). The association presented in Fig. 6C appears not to be strong enough for the model to appropriately capture age-related patch differences. Arguably, in datasets containing even more species (e.g. 50 – 100), the correlation might well be stronger, but in many situations, patch richness would also vary because of several factors beyond patch age. Therefore, its capacity to serve as a proxy of the latter would be compromised anyway. Consequently, using explicit independent data on patch age is the only generally recommendable strategy.

#### 3.2.4 Effect of changing parameter values and sample size

The performances of the main methods, under the different parameter scenarios, are summarized in Table 1. Confirming the previous results, only Hmsc M2 yielded satisfying results in all cases. All other methods generally detected spurious associations, often at quite high rates, sometimes exceeding 70%. Obviously, reducing sample size down to *N* = 300 reduced the general statistical power, and therefore the rate of spurious detections. In the case of species distribution model Hmsc M1, this sometimes made the percentage of errors fall to acceptable levels. However, this is just indicative of a lack of power, and would also come at the expense of higher type-II errors in actual datasets containing genuine species associations.

Decreasing the rate of patch disturbance *μ*, as expected, also decreases the over all amount of spurious associations detected (Table 1). This is simply because this weakens the metacommunity-dynamics driven patterns, in the absence of which all methods compared here perform correctly.

Interestingly, the effect of introducing some species immune to patch disturbance systematically reduces the rate of spurious detections for methods that do not control for patch richness (Fisher’s tests or Hmsc M0) but can have variable consequences for methods that do (fixed-fixed permutations and Hmsc M1). The consequences depend strongly on sample size. At low sample sizes, the main consequence is to reduce the average rate of perturbation, and thus to decrease the number of spurious detections, as statistical power is limiting. At large sizes, on the contrary, the presence of immune species increases the number of spurious detections, as it increases the heterogeneity in fastness among species, to which these methods are sensitive.

Our main overall conclusions are synthesized in Table 2.

**Table 2:** Are detection methods robust to patch disturbance? Summary of main findings. For each type of approach, we only mention methods tested in this study, but conclusions apply to other similar methods as well. Note that “Sim2” and “Sim9” denote the “fixed-equiprobable” and “fixed-fixed” algorithms, respectively.

| Type of approach | Example methods | Robust? | Type of bias |
| --- | --- | --- | --- |
| Direct independence tests | Fisher’s tests; Sim2; Hmsc M0 | No<sup>†</sup> | positive |
| Controlling for patch richness | Sim9; Hmsc M1; Hmsc M2’ | No* | positive/negative |
| Modelling effect of patch age | Hmsc M2 | Yes | none |

<sup>†</sup> works only if no patch disturbance \* works only if no patch disturbance, or all species similar (same fastness)

## 4 Discussion

Metacommunity dynamics matter. We demonstrated here how simple metacommunity dynamics can cause the detection of spurious species associations, even in null communities with no interactions. This occurs as soon as patch disturbances exist, i.e. catastrophic events that cause all species (or at least a subset of them) to be locally wiped out. The latter is an arguably common element in natural and exploited ecosystems (Pickett and White, 2013), and one often, though not always, included in metapopulation and metacommunity models (Leibold et al., 2004).

It is classically assumed that in the absence of any ecological interactions, habitat heterogeneity, or species differences in dispersal patterns, species would be distributed independently across patches. This “null” assumption underlies many interpretations of species co-occurrence patterns. Our analysis shows that this intuition is true for any given patch age (time since last disturbance), but it is true not overall. Patch age acts as a hidden structuring variable, the implications of which for co-occurrence patterns have been overlooked so far. Indeed, among the list of processes typically invoked as drivers which might cause departures from species independence, patch age does not normally appear (Gotelli, 2000; Ovaskainen and Abrego, 2020).

This result may seem counter-intuitive at first glance. Indeed, if species are initially independently distributed in a metacommunity, patch disturbances do not *per se* cause any association to appear: as they occur regardless of species composition, they preserve species independence, unlike interaction-driven extinctions would. Rather, it is the outcome of extinction/recolonization dynamics (Eq. 2) that inevitably causes, at steady-state, species to be more frequent in older patches. Since all species are more likely to occupy older patches, all appear positively associated overall.

The consequences of patch disturbance cannot simply be dealt with by controlling for patch differences in richness. If doing so, spurious associations, both negative and positive, also appear, as soon as species have different relative distribution profiles (fastnesses). These spurious associations have a different cause: they stem from the fact that slow and fast species are concentrated in different parts of the patch-age spectrum, which can be thought of as a form of niche differentiation. This differential distribution w.r.t. to patch age causes spurious negative association signals to appear. But the signature we report is more subtle than that (see Fig. 5). The fastest species (or species immune to patch disturbance, if any) also appear to be segregated from each other, as well as from all the other species. Reciprocally, the slowest species, being more concentrated than others in the older parts of the patch age spectrum, appear positively associated (and segregated from the others).

The generation of spurious associations (Type I error) is in itself problematic. But beyond that, in more realistic communities with actual species interactions, these spurious patterns would interfere with and confound the actual signals. For instance, actual signals of species interactions, e.g. an increase in species extinction rates with species number, i.e. a “neutral” form of competition (Hastings, 1987; Bell, 2001), could be hidden in the presence of patch disturbance: the negative association patterns would be cancelled, at least among pairs of slow species, by the positive signal caused by metacommunity dynamics alone. In such situations, false negatives (loss of power) will result. Furthermore, the power of detection is not homogeneously distributed across all species pairs: signals of competition would be weakened for slow species pairs, but reinforced for fast species pairs. Therefore, a diffuse type of competitive interaction with no discernible pattern (all species competitively identical) could turn into a structured pattern, with nonzero nestedness.

A possible solution to these issues consists of using species distribution models that explicitly incorporate patch age as a covariate (Fig. 6). However, this requires having independent data on patch ages, which is far from being universal in co-occurrence analyses. Whilst feasible for some types of systems (Ovaskainen et al., 2017), typically parasite or microbiome communities for which patches are hosts of known age, in many cases such data would be difficult or impossible to obtain. It may sometimes be possible to approximate patch-age from measurable environmental variables, when patch disturbance causes environmental modifications and patches follow a succession-like cycle after them. However, this approach could only be used for certain types of system, particularly since a sufficiently strong correlation would be required. Indeed, in our examples, using patch richness as a proxy of patch-age failed to remove the spurious associations, even though the two variables were reasonably well correlated (Fig. 6). In general, effectively controlling for patch age would almost certainly require genuine temporal data on patch life-cycles (Ovaskainen et al., 2017, Fig. 5c).

In general our work here underscores once more how inferring species interactions from snapshots of co-occurrence patterns can be as hazardous as it is appealing, and requires careful thought about null expectations (Blanchet et al., 2020; Molina and Stone, 2020; Münkemüller et al., 2020). We join the recent burst of cautionary articles in highlighting these concerns, and in any case, we encourage investigators to evaluate whether detected species association patterns could match those reported here, and so could stem from pure metacommunity dynamical effects.

## Acknowledgments

The authors thank Maxime Dubart for guidance on using Hmsc and for sharing data on Caribbean snail metacommunity parameter estimates. Colin Guétemme and Héloïse Villesèche participated in preliminary research on the topic.

## Data availability

All the code used to produce the results in this article is available on GitHub at https://github.com/nikcunniffe/MetacommunityDynamics.

## Supporting Information

The following additional information is available

- A. Mathematical supplements.
- B. Stochastic simulations.
- Supplementary Figure S1. Hmsc M2’: patch richness used as a proxy of patch age. (A) Species pairs for which a significant association is detected. (B) Same as (A) but with a set of immune species. Model Hmsc M2’ fails in a similar way as Hmsc M1 did (see Fig. 5C). For reference, Hmsc M2’ yields the following rates of errors: %+ = 6.7%, %- = 16%, %tot = 22.7% without immune species, and %+ = 8.8%, %- = 15%, %tot = 23.8% with immune species (see Table 1 for a comparison).

