## Supporting Information A for "Metacommunity dynamics and the detection of species associations in co-occurrence analyses: why patch disturbance matters"

### A. Mathematical supplements

**Vincent Calcagno, Nik J. Cunniffe, Frédéric M. Hamelin**

#### A1 General model

##### A1.1 Model and definitions

Patches undergo disturbances that lead to the extinction of all the species considered ($i = 1, 2, \dots, s$). The patches are disturbed at rate $\mu_x$, where $x$ is the age of a patch, i.e. the time since it was last disturbed. Let $X$ be the maximum age a patch can reach, i.e. patches are systematically disturbed at age $X$. In the special cases developed later, there is no such sharp limit, and $X$ just tends to infinity.

The model describes the changes in $p_{i,x,t}$, the mixed joint probability density of patch age $x$ (a continuous r.v.) and occupancy by species $i$ (a discrete r.v.) at time $t$. The marginal probability of occupancy by species $i$ is

$$p_{i,\bullet,t} = \int_0^X p_{i,x,t} \mathrm{d}x \,.$$

Similarly, the marginal probability density of patch age $x$ at time $t$ is denoted as $p_{\bullet,x,t}$ (see Section A3 for a more explicit definition), and sums to unity as any p.d.f.:

$$\int_0^X p_{\bullet,x,t} \mathrm{d}x = 1 \,.$$

Lastly, the probability that a patch of age $x$ is occupied by species $i$ at time $t$ is denoted as:

$$p_{i|x,t} = \frac{p_{i,x,t}}{p_{\bullet,x,t}} .$$

The general model can be expressed as the following partial differential equation (repeating Eq. 1 in the main text):

$$\frac{\partial p_{i,x,t}}{\partial x} + \frac{\partial p_{i,x,t}}{\partial t} = -(\mu_x + e_i)\, p_{i,x,t} + (c_i p_{i,\bullet,t} + m_i)\,(p_{\bullet,x,t} - p_{i,x,t}) \,. \qquad \text{(A-1)}$$

Since all patches are empty following a disturbance, $p_{i,0,t} = 0$ for all $i = 1, 2, \ldots, s$, and for all $t \geq 0$. If there is a maximum patch age $X$, $p_{i,x,t} = 0$ for all $x > X$. Otherwise, $\lim_{x\to+\infty} p_{i,x,t} = 0$.

At steady-state, we can drop the subscript $t$, and Eq. A-1 becomes:

$$\frac{\mathrm{d}p_{i,x}}{\mathrm{d}x} = -(\mu_x + e_i)\, p_{i,x} + (c_i p_{i,\bullet} + m_i)\,(p_{\bullet,x} - p_{i,x}) \,, \qquad \text{(A-2)}$$

where $p_{\bullet,x}$ is the stationary probability density of the age $x$ of a patch.

Table A1 lists the model parameters/variables and their definitions.

#### A1.2 Steady state distribution of patch age

The stationary distribution of patch age satisfies

$$\frac{\mathrm{d}p_{\bullet,x}}{\mathrm{d}x} = -\mu_x p_{\bullet,x} \,.$$

Therefore, $p_{\bullet,x}$ can be expressed as:

$$p_{\bullet,x} = p_{\bullet,0} \exp\left(-\int_0^x \mu_y \mathrm{d}y\right) , \qquad \text{(A-3)}$$

where $p_{\bullet,0}$ is an implicit factor such that $\int_0^X p_{\bullet,x}\mathrm{d}x = 1$, since $p_{\bullet,x}$ is the probability density function of the host age $x$. The function $p_{\bullet,x}$ is decreasing with respect to $x$. In the special case where $\mu_x = \mu$ (a constant) and $X$ is infinite, it is simply an exponential distribution with rate $\mu$. In general, depending on $\mu_x$, $p_{\bullet,x}$ can take various shapes, including for instance uniform or Weibull distributions.

| Parameter | Meaning |
|---|---|
| $s$ | number of species considered; species are indexed with $i, j = 1, 2, ..., s$ |
| $c_i$ | colonization rate of species $i$ (per occupied patch) |
| $m_i$ | immigration rate from outside the metacommunity of species $i$ |
| $e_i$ | local extinction rate of species $i$ |
| $\mu_x$ | catastrophic disturbance rate of a patch of age $x$ (noted $\mu$ if constant) |
| $X$ | maximum patch age (if any) |
| $N$ | total number of sites (patches) in the co-occurrence matrix considered |
| **Variable** | **Meaning** |
| $t$ | time |
| $x$ | age of a patch, i.e. the time since the last catastrophic disturbance event |
| $F_{i,t}$ | force of colonization/immigration of species $i$ at time $t$ |
| $F_i$ | steady-state force of colonization/immigration of species $i$ |
| $p_{i,x,t}$ | fraction of patches that have age $x$ and are occupied by species $i$ at time $t$ |
| $p_{\bullet,x,t}$ | fraction of patches that have age $x$ at time $t$ |
| $p_{i,\bullet,t}$ | overall occupancy of species $i$, i.e. the fraction of patches it occupies |
| $p_{i\vert x,t}$ | fraction of patches of age $x$ that are occupied by spp. $i$: $p_{i\vert x,t} = p_{i,x,t}/p_{\bullet,x,t}$ |
| $p_{i,x}$ | steady-state fraction of patches that have age $x$ and are occupied by spp. $i$ |
| $p_{\bullet,x}$ | steady-state fraction of patches that have age $x$ |
| $p_{i,\bullet}$ | overall occupancy of species $i$ at steady-state |
| $p^*_{i,\bullet}$ | overall occupancy of species $i$ after permutations in the co-occurrence matrix |
| $p_{i\vert x}$ | fraction of patches of age $x$ that are occupied by species $i$: $p_{i\vert x} = p_{i,x}/p_{\bullet,x}$ |
| $\pi_{i/x}$ | relative distribution profile of species $i$: $\pi_{i/x} = p_{i\vert x}/p_{i,\bullet}$ |
| $\pi^*_{i/x}$ | relative distribution profile of species $i$ after permutations in the matrix |
| $\pi^{-1}_{i/z}$ | inverse function of $\pi_{i/x}$ |
| $p_{i,z}$ | the probability density function of $\pi_{i/x}$ |
| $q_{\varnothing,t,x}$ | fraction of patches that have age $x$ and are unoccupied at time $t$ |
| $q_{i,t,x}$ | fraction of patches that have age $x$ and are occupied by a single species $i$ |
| $q_{\{i,j\},t,x}$ | fraction of patches that have age $x$ and are occupied by both species $i$ and $j$ |
| $q_{i,t,\bullet}$ | overall fraction of patches occupied by species $i$ only at time $t$ |
| $q_{\{i,j\},t,\bullet}$ | overall fraction of patches occupied by both species $i$ and $j$ at time $t$ |
| $q_{\varnothing,x}$ | steady-state fraction of patches that have age $x$ and are unoccupied |
| $q_{i,x}$ | steady-state fraction of patches that have age $x$ and are singly occupied |
| $q_{\{i,j\},x}$ | steady-state fraction of patches that have age $x$ and are doubly occupied |
| $q_{i,j,\bullet}$ | overall fraction of patches occupied by both species $i$ and $j$ at steady-state |
| $q^*_{i,j,\bullet}$ | overall fraction of co-occurrences of species $i$ and $j$ after permutations |
| $C_{i,j}$ | partial C-score between two species: $C_{i,j} = N^2(p_{i,\bullet} - q_{i,j,\bullet})(p_{j,\bullet} - q_{i,j,\bullet})$ |
| $w_i$ | relative occupancy of species $i$ in the matrix: $w_i = p_{i,\bullet}/\sum_{k=1}^{s} p_{k,\bullet}$ |
| $\hat{\pi}_x$ | weighted-average of the species distribution profiles: $\hat{\pi}_x = \sum_{k=1}^{s} w_k \pi_{k/x}$ |

Table A1: Model variables and parameters.

#### A1.3 Occupancy conditional on patch age

From Eq. A-2 and A-3 and the rule of differentiation of a ratio ($p_{i,x}/p_{\bullet,x}$), the steady-state probability of occupancy conditioned to patch age ($p_{i|x}$) satisfies:

$$\frac{\mathrm{d}p_{i|x}}{\mathrm{d}x} = -e_i p_{i|x} + (c_i p_{i,\bullet} + m_i)\left(1 - p_{i|x}\right),$$

with $p_{i|0} = 0$. This can be solved as

$$p_{i|x} = \frac{c_i p_{i,\bullet} + m_i}{c_i p_{i,\bullet} + m_i + e_i}\left(1 - \exp(-(c_i p_{i,\bullet} + m_i + e_i)x)\right). \qquad \text{(A-4)}$$

#### A1.4 Overall steady state occupancy

The steady state occupancy of a species is defined implicitly by

$$p_{i,\bullet} = \int_0^X p_{i|x} p_{\bullet,x} \mathrm{d}x, \qquad \text{(A-5)}$$

with $p_{i|x}$ given in Eq. A-4.

In general, this admits no explicit solution. However, we can solve for $p_{i,\bullet}$ using a simple iterative algorithm:

1. Set $p_{i,\bullet}$ to some non-zero initial value, e.g. $\frac{1}{2}$;
2. Update its value using eq. A-5;
3. Repeat 2 until the value of $p_{i,\bullet}$ no longer changes (fixed point).

#### A1.5 Relative distribution profiles

The relative distribution profile of species $i$ is defined as in Eq. 2 in the main text:

$$\pi_{i/x} = \frac{p_{i|x}}{p_{i,\bullet}} = \frac{1}{p_{i,\bullet}} \frac{c_i p_{i,\bullet} + m_i}{c_i p_{i,\bullet} + m_i + e_i}\left(1 - \exp\left(-(c_i p_{i,\bullet} + m_i + e_i)x\right)\right). \qquad \text{(A-6)}$$

We note that the mean value of the profile is one:

$$\mathrm{E}(\pi_{i/x}) = \int_0^X \pi_{i/y} p_{\bullet,y} \mathrm{d}y = \int_0^X \frac{p_{i,y}}{p_{\bullet,y} p_{i,\bullet}} p_{\bullet,y} \mathrm{d}y = 1. \qquad \text{(A-7)}$$

Let

$$A_i = \frac{1}{p_{i,\bullet}} \frac{c_i p_{i,\bullet} + m_i}{c_i p_{i,\bullet} + m_i + e_i}\,, \quad \text{and} \quad R_i = c_i p_{i,\bullet} + m_i + e_i\,.$$

We have:

$$\pi_{i/x} = A_i\left[1 - \exp\left(-R_i x\right)\right]\,.$$

We note that $\pi_{i,0} = 0$ for all $i$, and that $\lim_{x \to X} \pi_{i/x} = A_i(1 - \exp(-R_i X))$. The latter is between 1 and $A_i$, since on the one hand $\pi_{i/x}$ is strictly increasing w.r.t. $x$, and $\mathrm{E}(\pi_{i/x}) = 1$, and on the other hand $\lim_{x \to \infty} \pi_{i/x} = A_i$.

To obtain the distribution (probability density) of $\pi_{i/x}$ values, we first compute the inverse function $\pi^{-1}_{i/z}$, that returns the patch age for which a particular value $z$ of $\pi_{i/x}$ is obtained. From the above expression of $\pi_{i/x}$ we get:

$$\pi^{-1}_{i/z} = \frac{1}{R_i} \ln\left[\frac{A_i}{A_i - z}\right]\,.$$

It then follows that the probability density function of $\pi_{i/x}$ is:

$$p_{i,z} = \frac{\mathrm{d}\pi^{-1}_{i/z}}{\mathrm{d}z} p_{\bullet,\pi^{-1}_{i,z}} = \frac{p_{\bullet,\pi^{-1}_{i,z}}}{R_i(A_i - z)}\,,$$

defined on the interval $0 < z < A_i(1 - \exp(-R_i X))$. This expression was used to draw the distribution of $\pi_{i/x}$ values in the inset of Figure 2 in the main text. The variance of the above distribution is a metric of species "fastness" (see main text).

We also note that since $\pi_{i/x}$ and $\pi_{j/x}$ are increasing functions of $x$ (Eq. A-6), Harris' inequality applies:

$$\int_0^X p_{\bullet,x} \pi_{i/x} \pi_{j/x} \mathrm{d}x \geq \int_0^X p_{\bullet,x} \pi_{i/x} \mathrm{d}x \int_0^X p_{\bullet,x} \pi_{j/x} \mathrm{d}x = 1\,, \qquad \text{(A-8)}$$

or equivalently $\mathrm{Cov}(\pi_{i/x}, \pi_{j/x}) \geq 0$.

Lastly, we remark that the variance of the average community profile (see Eq. 9 in the main text) can also be expressed as:

$$\mathrm{Var}(\hat{\pi}) = \sum_{k=1}^{S} \sum_{\ell=1}^{S} w_k w_\ell \,\mathrm{Cov}(\pi_{k/x}, \pi_{\ell/x})\,.$$

#### A1.6 The relative distribution profiles cross exactly once

The following lemma will be used in the theorem of the next section.

**Lemma.** *In the general metacommunity model A-1, for any pair of species* $i, j$*, the relative distribution profiles* $\pi_{i/x}$ *and* $\pi_{j/x}$ *cross exactly once beyond the initial point* $(x = 0)$*, unless* $\pi_{i/x} = \pi_{j/x}$ *for all* $x$*.*

*Proof.* Let

$$\delta_x = \pi_{i/x} - \pi_{j/x} = A_i \left[1 - \exp\left(-R_i x\right)\right] - A_j \left[1 - \exp\left(-R_j x\right)\right] .$$

Differentiating with respect to $x$,

$$\frac{\mathrm{d}\delta_x}{\mathrm{d}x} = A_i R_i \exp\left(-R_i x\right) - A_j R_j \exp\left(-R_j x\right) ,$$

with $\delta_0 = 0$. Let us show that $\delta_x$ has a single optimum (maximum or minimum). Let $x^*$ be such that $\delta'_{x^*} = 0$, where the prime denotes the slope of $\delta_x$. We find a unique possible such $x^*$:

$$x^* = \frac{\log\left(\frac{A_i R_i}{A_j R_j}\right)}{R_i - R_j} .$$

Therefore, regardless the initial sign of $\delta'_x$, $\delta_x$ cannot change sign more than once (for some $x > x^*$). Since

$$\mathrm{E}(\delta_x) = \int_0^X p_{\bullet,x} \delta_x \mathrm{d}x = \int_0^X p_{\bullet,x} (\pi_{i,x} - \pi_{i,x}) = 0 ,$$

$\delta_x$ must change sign at least once. Hence, $\delta_x$ changes sign exactly once. That means that any pair of profiles $\pi_{i/x}, \pi_{j/x}$ cross exactly once beyond the initial point $(x = 0)$. □

#### A1.7 Variances and initial slopes of the relative distribution profiles

**Theorem.** *In the general metacommunity model A-1, for any pair of species* $i, j$*, the inequality* $\mathrm{Var}(\pi_{i/x}) < \mathrm{Var}(\pi_{j/x})$ *is equivalent to* $\pi'_{i/0} > \pi'_{j/0}$*, meaning that the variance of the profile is entirely determined by the initial slope of the profile.*

*Proof.* The inequality $\mathrm{Var}(\pi_{i/x}) < \mathrm{Var}(\pi_{j/x})$ is equivalent to

$$\int_0^X \pi_{i/x}^2 p_{\bullet,x} \mathrm{d}x < \int_0^X \pi_{j/x}^2 p_{\bullet,x} \mathrm{d}x\,.$$

This inequality can equivalently be expressed as:

$$\int_0^X (\pi_{i/x}^2 - \pi_{j/x}^2) p_{\bullet,x} \mathrm{d}x = \int_0^X (\pi_{i/x} + \pi_{j|x})(\pi_{i/x} - \pi_{j/x}) p_{\bullet,x} \mathrm{d}x = \int_0^X (\pi_{i/x} + \pi_{j/x}) \delta_x p_{\bullet,x} \mathrm{d}x < 0\,.$$

Since $(\pi_{i/x} + \pi_{j/x})$ is increasing w.r.t. $x$, $\mathrm{E}(\delta_x) = 0$, and using the preceding lemma, it is necessary and sufficient that $\delta_x$ is positive on the interval $(0, x^*)$ for the above inequality to be satisfied. This condition is satisfied if and only if $\delta_0' = A_i R_i - A_j R_j > 0$. Therefore, the above inequality is equivalent to $A_i R_i > A_j R_j$. For $k = i, j$,

$$A_k R_k = c_k + \frac{m_k}{p_{k,\bullet}} = \pi_{k/0}'\,,$$

where the prime denotes differentiation w.r.t. $x$. Hence the equivalence:

$$\mathrm{Var}(\pi_{i/x}) < \mathrm{Var}(\pi_{j/x}) \Longleftrightarrow \pi_{i/0}' > \pi_{j/0}'\,,$$

□

**Note:** this equivalence between initial slope and variance holds for any species, but does not hold for the average profile of several species. Therefore the initial slope of the average relative distribution profile cannot be taken as a proxy for $\mathrm{Var}(\hat{\pi})$. It is therefore not a good proxy of average fastness.

#### A2 Link with classical metapopulation models

The model (Eq. A-1) generalizes the classical mainland-island and Levins metapopulation models, which are characterized by $\mu(x) = \mu$ (a constant) for all $x$, and $X \to +\infty$. To show the connection, we integrate both sides of Eq. A-1 over $x$ on $[0, +\infty)$. The l.h.s. simplifies to

$$\left(\lim_{x\to+\infty} p_{i,x,t} - p_{i,0,t}\right) + \frac{\mathrm{d}p_{i,\bullet,t}}{\mathrm{d}t} = \frac{\mathrm{d}p_{i,\bullet,t}}{\mathrm{d}t}\,,$$

which yields

$$\frac{\mathrm{d}p_{i,\bullet,t}}{\mathrm{d}t} = (c_i p_{i,\bullet,t} + m_i)(1 - p_{i,\bullet,t}) - (\mu + e_i) p_{i,\bullet,t} \,. \tag{A-9}$$

We recognize a classical metapopulation model. The special cases $c_i = 0$ and $m_i = 0$ correspond to the mainland-island and Levins models, respectively.

#### A2.1 Steady-state occupancy in classical models

At steady-state, Eq. A-9 becomes:

$$0 = (c_i p_{i,\bullet} + m_i)(1 - p_{i,\bullet}) - (\mu + e_i) p_{i,\bullet} \,. \tag{A-10}$$

Solving for $p_{i,\bullet}$ in Eq. A-10 yields two real roots. One can easily check that only the largest root is positive. The biologically relevant equilibrium is therefore

$$p_{i,\bullet} = \frac{c_i - (e_i + \mu + m_i) + \sqrt{m_i^2 + 2(e_i + c_i + \mu) m_i + (e_i - c_i + \mu)^2}}{2 c_i} \,, \tag{A-11}$$

which requires $c_i > 0$.

**Mainland-island model.** Assuming $c_i = 0$, solving for $p_{i,\bullet}$ in Eq. A-10 yields

$$p_{i,\bullet} = \frac{m_i}{m_i + e_i + \mu} \,. \tag{A-12}$$

**Levins model.** Assuming $m_i = 0$, Eq. A-11 simplifies to

$$p_{i,\bullet} = 1 - \frac{e_i + \mu}{c_i} \,, \tag{A-13}$$

provided $c_i > e_i + \mu$. Otherwise $p_{i,\bullet} = 0$.

#### A2.2 Expressions of the variance/covariance of relative profiles

The overall fractions of patches occupied by species $i$ only, species $j$ only, and both species $i$ and species $j$, are $q_{i,t,\bullet}$, $q_{j,t,\bullet}$, and $q_{\{i,j\},t,\bullet}$, respectively. The fractions of patches occupied by species $i$ and species $j$ are $p_{i,t,\bullet} = q_{i,t,\bullet} + q_{\{i,j\},t,\bullet}$ and $p_{j,t,\bullet} =$

$q_{j,t,\bullet} + q_{\{i,j\},t,\bullet}$, respectively. Integrating both sides of Eq. A-14 w.r.t. $x$,

$$\begin{aligned}
\frac{\mathrm{d}q_{i,t,\bullet}}{\mathrm{d}t} &= F_{i,t}(1 - q_{i,t,\bullet} - q_{j,t,\bullet} - q_{\{i,j\},t,\bullet}) - e_i q_{i,t,\bullet} - \mu q_{i,t,\bullet} + e_j q_{\{i,j\},t,\bullet} - F_{j,t} q_{i,t,\bullet}\,,\\
\frac{\mathrm{d}q_{j,t,\bullet}}{\mathrm{d}t} &= F_{j,t}(1 - q_{i,t,\bullet} - q_{j,t,\bullet} - q_{\{i,j\},t,\bullet}) - e_j q_{j,t,\bullet} - \mu q_{j,t,\bullet} + e_i q_{\{i,j\},t,\bullet} - F_{i,t} q_{j,t,\bullet}\,,\\
\frac{\mathrm{d}q_{\{i,j\},t,\bullet}}{\mathrm{d}t} &= F_{j,t} q_{i,t,\bullet} + F_{i,t} q_{j,t,\bullet} - (e_i + e_j + \mu) q_{\{i,j\},t,\bullet}\,.
\end{aligned}$$

The above system of equations can equivalently be expressed as:

$$\begin{aligned}
\frac{\mathrm{d}p_{i,t,\bullet}}{\mathrm{d}t} &= F_{i,t}(1 - p_{i,t,\bullet}) - (e_i + \mu) p_{i,t,\bullet}\,,\\
\frac{\mathrm{d}p_{j,t,\bullet}}{\mathrm{d}t} &= F_{j,t}(1 - p_{j,t,\bullet}) - (e_j + \mu) p_{j,t,\bullet}\,,\\
\frac{\mathrm{d}q_{\{i,j\},\bullet}}{\mathrm{d}t} &= F_{j,t}(p_{i,t,\bullet} - q_{\{i,j\},t,\bullet}) + F_{j,t}(p_{j,t,\bullet} - q_{\{i,j\},t,\bullet}) - (e_i + e_j + \mu) q_{\{i,j\},\bullet}\,.
\end{aligned}$$

At steady-state, we can drop the $t$ subscripts: for $k = i, j$,

$$p_{k,\bullet} = \frac{F_k}{F_k + e_k + \mu}\,, \quad \text{and} \quad q_{\{i,j\},\bullet} = \frac{F_j p_{i,\bullet} + F_i p_{j,\bullet}}{F_j + F_i + e_i + e_j + \mu} = p_{i,\bullet} p_{j,\bullet} \frac{\frac{F_j}{p_{j,\bullet}} + \frac{F_i}{p_{i,\bullet}}}{F_i + F_j + e_i + e_j + \mu}\,.$$

Combining both equations,

$$q_{\{i,j\},\bullet} = p_{i,\bullet} p_{j,\bullet} \frac{F_i + F_j + e_i + e_j + 2\mu}{F_i + F_j + e_i + e_j + \mu} = p_{i,\bullet} p_{j,\bullet} \left(1 + \frac{\mu}{F_i + F_j + e_i + e_j + \mu}\right).$$

Therefore (see Eq. 5 in the main text),

$$\mathrm{Cov}(\pi_{i/x}, \pi_{j/x}) = \frac{\mu}{F_i + F_j + e_i + e_j + \mu}\,.$$

Using the fact that the force of colonization/immigration of species $i$ (Eq. A-15) is $F_i = c_i(q_{i,\bullet} + q_{\{i,j\},\bullet}) + m_i = c_i p_{i,\bullet} + m_i$,

$$\mathrm{Cov}(\pi_{i/x}, \pi_{j/x}) = \frac{\mu}{c_i p_{i,\bullet} + m_i + c_j p_{j,\bullet} + m_j + e_i + e_j + \mu}\,.$$

Last, using the expression of $p_{i,\bullet}$ from Eq. A-11, we obtain:

$$\mathrm{Cov}(\pi_{i/x}, \pi_{j/x}) = \frac{2\mu}{\sum_{k\in\{i,j\}} \left(m_k + e_k + c_k + \sqrt{m_k^2 + 2(e_k + c_k + \mu) m_k + (e_k - c_k + \mu)^2}\right)}\,.$$

As a special case, for any species $i$:

$$\mathrm{Var}(\pi_{i,x}) = \mathrm{Cov}(\pi_{i/x}, \pi_{i/x}) = \frac{\mu}{m_i + e_i + c_i + \sqrt{m_i^2 + 2(e_i + c_i + \mu)m_i + (e_i - c_i + \mu)^2}}\,.$$

**Mainland-island model.** Assuming $c_i = 0$ and rearranging yields:

$$\mathrm{Cov}(\pi_{i/x}, \pi_{j/x}) = \frac{\mu}{m_i + m_j + e_i + e_j + \mu}\,, \quad \text{and} \quad \mathrm{Var}(\pi_{i/x}) = \frac{\mu}{2(e_i + m_i) + \mu}\,.$$

**Levins model.** Assuming $m_i = 0$, using and rearranging yields:

$$\mathrm{Cov}(\pi_{i/x}, \pi_{j/x}) = \frac{\mu}{c_i + c_j - \mu}\,, \quad \text{and} \quad \mathrm{Var}(\pi_{i/x}) = \frac{\mu}{2c_i - \mu}\,.$$

#### A2.3 Parameter conditions to have identical relative profiles

We define as similar species that have the same relative distribution profile: for any pair of similar species $i, j$, $\pi_{i/x} = \pi_{j/x}$ for all $x$. The latter equality is equivalent to $p_{i|x} = \kappa p_{j|x}$, which is again equivalent to $p_{i,x} = \kappa p_{j,x}$, with $\kappa = p_{i,\bullet}/p_{j,\bullet}$.

Using Eq. A-4, this means that the following pair of equations must be satisfied:

$$\begin{aligned} c_i p_{i,\bullet} + m_i + e_i &= c_j p_{j,\bullet} + m_j + e_j\,, \\ c_i p_{i,\bullet} + m_i &= \kappa(c_j p_{j,\bullet} + m_j)\,. \end{aligned}$$

**Mainland-island model.** Assuming $c_i = 0$, using Eq. A-12, and rearranging yields:

$$\begin{aligned} m_i &= \kappa m_j\,, \\ e_i &= (1-\kappa)m_j + e_j\,. \end{aligned}$$

In the mainland-island model, similar species may differ in both extinction and immigration rates, provided they respect the above relationships.

**Levins model.** Assuming $m_i = 0$, using Eq. A-13, and rearranging yields:

$$\begin{aligned} c_i &= c_j\,, \\ e_i &= (1-\kappa)(c_j - \mu) + \kappa e_j\,. \end{aligned}$$

In the Levins model, similar species must have equal colonization rates, but their extinction rates may differ provided they respect the above relationship.

#### A3 Independence of co-occurrences within age classes

In this section, we keep track of co-occurrences between two non-interacting species. We will show that the probability that a patch of age $x$ is occupied by both species is equal to the product of the probabilities that a patch of age $x$ is occupied by each species irrespective of the other species. The demonstration is inspired from and extends earlier studies in epidemiology (Kucharski and Gog, 2012; Hamelin et al., 2019).

We consider the following model, which looks into the general metacommunity model A-1 into more detail for any pair of species in the set of $s$ species considered. These species are indexed by $i = 1, 2$ without loss of generality. Let $q_{\varnothing,t,x}$, $q_{i,t,x}$, and $q_{\{1,2\},t,x}$ be the fractions of patches unoccupied, occupied by a single species ($i = 1, 2$), and occupied by both species, respectively. For $x > 0$,

$$\begin{aligned}
\frac{\partial q_{\varnothing,t,x}}{\partial t} + \frac{\partial q_{\varnothing,t,x}}{\partial x} &= -(F_{1,t} + F_{2,t} + \mu_x) q_{\varnothing,t,x} + e_1 q_{1,t,x} + e_2 q_{2,t,x}\,, \\
\frac{\partial q_{1,t,x}}{\partial t} + \frac{\partial q_{1,t,x}}{\partial x} &= F_{1,t} q_{\varnothing,t,x} - (F_{2,t} + \mu_x + e_1) q_{1,t,x} + e_2 q_{\{1,2\},t,x}\,, \\
\frac{\partial q_{2,t,x}}{\partial t} + \frac{\partial q_{2,t,x}}{\partial x} &= F_{2,t} q_{\varnothing,t,x} - (F_{1,t} + \mu_x + e_2) q_{2,t,x} + e_1 q_{\{1,2\},t,x}\,, \\
\frac{\partial q_{\{1,2\},t,x}}{\partial t} + \frac{\partial q_{\{1,2\},t,x}}{\partial x} &= F_{2,t} q_{1,t,x} + F_{1,x} q_{2,t,x} - (\mu_x + e_1 + e_2) q_{\{1,2\},t,x}\,,
\end{aligned} \qquad \text{(A-14)}$$

and, for $x = 0$:

$$\begin{aligned}
q_{\varnothing,t,0} &= \int_0^X \mu_x (q_{\varnothing,t,x} + q_{1,t,x} + q_{2,t,x} + q_{\{1,2\},t,x}) \mathrm{d}x + q_{\varnothing,t,X} + q_{1,t,X} + q_{2,t,X} + q_{\{1,2\},t,X}\,, \\
q_{1,t,0} &= 0\,, \quad q_{2,t,0} = 0\,, \quad q_{\{1,2\},t,0} = 0\,.
\end{aligned}$$

The force of colonization/immigration of species $i = 1, 2$, $F_{i,t}$, can for instance take the form:

$$F_{i,t} = c_i \int_0^X (q_{1,t,x} + q_{\{1,2\},t,x}) \mathrm{d}x + m_i\,.$$

We set $p_{\bullet,t,x} = q_{\varnothing,t,x} + q_{1,t,x} + q_{2,t,x} + q_{\{1,2\},t,x}$. We have, for $x > 0$,

$$\frac{\partial p_{\bullet,t,x}}{\partial t} + \frac{\partial p_{\bullet,t,x}}{\partial x} = -\mu_x p_{\bullet,t,x}\,,$$

and, for $x = 0$,

$$p_{\bullet,t,0} = \int_0^X \mu_x p_{\bullet,t,x} \mathrm{d}x + p_{\bullet,t,X}\,.$$

**Steady-state analysis.** At steady state, the state variables do not depend on time $t$, and the model simplifies to (keeping the same notations for convenience):

$$\begin{aligned}
\frac{\mathrm{d}q_{\varnothing,x}}{\mathrm{d}x} &= -(F_1 + F_2 + \mu_x) q_{\varnothing,x} + e_1 q_{1,x} + e_2 q_{2,x}\,, \\
\frac{\mathrm{d}q_{1,x}}{\mathrm{d}x} &= F_1 q_{\varnothing,x} - (F_2 + \mu_x + e_1) q_{1,x} + e_2 q_{\{1,2\},x}\,, \\
\frac{\mathrm{d}q_{2,x}}{\mathrm{d}x} &= F_2 q_{\varnothing,x} - (F_1 + \mu_x + e_2) q_{2,x} + e_1 q_{\{1,2\},x}\,, \\
\frac{\mathrm{d}q_{\{1,2\},x}}{\mathrm{d}x} &= F_2 q_{1,x} + F_1 q_{2,x} - (\mu_x + e_1 + e_2) q_{\{1,2\}}\,,
\end{aligned}$$

with initial conditions:

$$\begin{aligned}
q_{\varnothing,0} &= \int_0^X \mu_x (q_{\varnothing,x} + q_{1,x} + q_{2,x} + q_{\{1,2\},x}) \mathrm{d}x + q_{\varnothing,X} + q_{1,X} + q_{2,X} + q_{\{1,2\},X}\,, \\
q_{1,0} &= 0\,, \quad q_{2,0} = 0\,, \quad q_{\{1,2\},0} = 0\,.
\end{aligned}$$

The force of colonization/immigration of species $i = 1,2$, $F_i$, can for instance take the form:

$$F_i = c_i \int_0^X (q_{1,x} + q_{\{1,2\},x}) \mathrm{d}x + m_i\,. \tag{A-15}$$

We set $p_{\bullet,x} = q_{\varnothing,x} + q_{1,x} + q_{2,x} + q_{\{1,2\},x}$. The distribution $p_{\bullet,x}$ has been expressed in Eq. A-3.

Next we define the probabilities for a patch to be unoccupied, occupied by a single species ($i = 1,2$), and occupied by both species, given patch age $x$:

$$q_{\varnothing|x} = \frac{q_{\varnothing,x}}{p_{\bullet,x}}\,, \quad q_{1|x} = \frac{q_{1,x}}{p_{\bullet,x}}\,, \quad q_{2|x} = \frac{q_{2,x}}{p_{\bullet,x}}\,, \quad q_{\{1,2\}|x} = \frac{q_{\{1,2\},x}}{p_{\bullet,x}}\,.$$

Note that $q_{\varnothing|x}+q_{1|x}+q_{2|x}+q_{\{1,2\}|x}=1$. Using the fact that

$$q'_{\varnothing|x}=\left(\frac{q_{\varnothing,x}}{p_{\bullet,x}}\right)'=\frac{q'_{\varnothing,x}}{p_{\bullet,x}}-q_{\varnothing|x}\frac{p'_{\bullet,x}}{p_{\bullet,x}}=\frac{q'_{\varnothing,x}}{p_{\bullet,x}}+\mu_x q_{\varnothing|x}\,,$$

and similarly for other conditional probabilities, we obtain:

$$\begin{aligned}
\frac{\mathrm{d}q_{\varnothing|x}}{\mathrm{d}x} &= -(F_1+F_2)q_{\varnothing|x}+e_1q_{1|x}+e_2q_{2|x}\,,\\
\frac{\mathrm{d}q_{1|x}}{\mathrm{d}x} &= F_1q_{\varnothing|x}-(F_2+e_1)q_{1|x}+e_2q_{\{1,2\}|x}\,,\\
\frac{\mathrm{d}q_{2|x}}{\mathrm{d}x} &= F_2q_{\varnothing|x}-(F_1+e_2)q_{2|x}+e_1q_{\{1,2\}|x}\,,\\
\frac{\mathrm{d}q_{\{1,2\}|x}}{\mathrm{d}x} &= F_2q_{1|x}+F_1q_{2|x}-(e_1+e_2)q_{\{1,2\}|x}\,,
\end{aligned}$$

with initial conditions

$$\begin{aligned}
q_{\varnothing|0} &= \frac{q_{\varnothing,0}}{p_{\bullet,0}}=1\,,\\
q_{1|0} &= 0\,,\quad q_{2|0}=0\,,\quad q_{\{1,2\}|0}=0\,.
\end{aligned}$$

Let

$$p_{1|x}=q_{1|x}+q_{\{1,2\}|x}\,,\quad p_{2|x}=q_{2|x}+q_{\{1,2\}|x}\,,\quad \Delta_x=q_{\{1,2\}|x}-p_{1|x}p_{2|x}\,.$$

We have

$$\begin{aligned}
\frac{\mathrm{d}p_{1|x}}{\mathrm{d}x} &= F_1(q_{\varnothing|x}+q_{2|x})-e_1(q_{1|x}+q_{\{1,2\}|x})\,,\\
&= F_1(1-p_{1|x})-e_1p_{1|x}\,,
\end{aligned}$$

and similarly for the derivative of $p_{2|x}$ w.r.t. $x$. Thus

$$\frac{\mathrm{d}(p_{1|x}p_{2|x})}{\mathrm{d}x}=F_1(1-p_{1|x})p_{2|x}+F_2(1-p_{2|x})p_{1|x}-(e_1+e_2)p_{1|x}p_{2|x}\,.$$

Using

$$\begin{aligned}
\frac{\mathrm{d}q_{\{1,2\}|x}}{\mathrm{d}x} &= F_1(p_{2|x}-q_{\{1,2\}|x})+F_2(p_{1|x}-q_{\{1,2\}|x})-(e_1+e_2)q_{\{1,2\}|x}\,,\\
\frac{\mathrm{d}(p_{1|x}p_{2|x})}{\mathrm{d}x} &= F_1(p_{2|x}-p_{1|x}p_{2|x})+F_2(p_{1|x}-p_{1|x}p_{2|x})-(e_1+e_2)p_{1|x}p_{2|x}\,,
\end{aligned}$$

we end up with:

$$\frac{\mathrm{d}\Delta_x}{\mathrm{d}x} = -(F_1 + F_2 + e_1 + e_2)\Delta_x\,, \quad \Delta_0 = 0\,.$$

Therefore, $\Delta_x = 0$ for all $x$. Hence, for all $x \geq 0$,

$$q_{\{1,2\}|x} = p_{1|x} p_{2|x}\,.$$

As a consequence,

$$q_{1,2,\bullet} = \int_0^X q_{\{1,2\}|x} p_{\bullet,x} \mathrm{d}x = \int_0^X p_{1|x} p_{2|x} p_{\bullet,x} \mathrm{d}x\,.$$

### A4 The case of immune species

In this section, we consider species that are immune to disturbance events (hereafter immune species) separately from vulnerable (non-immune) species.

For immune species, the boundary conditions of the general model A-1 have to be updated in the following way: for all immune species indexed by $i$, and for all $t \geq 0$,

$$p_{i,0,t} = \int_0^X \mu_x p_{i,x,t} \mathrm{d}x + p_{i,X,t}\,.$$

Since species have nonzero individual extinction rates (for all $i$, $e_i > 0$), we have $\lim_{X \to +\infty} p_{i,X,t} = 0$.

The patches that have age zero at time $t$ are those that are disturbed at time $t$:

$$p_{\bullet,0,t} = \int_0^X \mu_x p_{\bullet,x,t} \mathrm{d}x + p_{\bullet,X,t}\,.$$

Integrating both sides of Eq. A-1 w.r.t. $x$,

$$(p_{i,X,t} - p_{i,0,t}) + \frac{\mathrm{d}p_{i,\bullet,t}}{\mathrm{d}t} = -\int_0^X \mu_x p_{i,x,t} \mathrm{d}x - e_i p_{i,\bullet,t} + (c_i p_{i,\bullet,t} + m_i)(1 - p_{i,\bullet,t})\,,$$

which yields

$$\frac{\mathrm{d}p_{i,\bullet,t}}{\mathrm{d}t} = (c_i p_{i,\bullet,t} + m_i)(1 - p_{i,\bullet,t}) - e_i p_{i,\bullet,t}\,.$$

At steady-state, we omit the $t$ subscripts:

$$0 = (c_i p_{i,\bullet} + m_i)(1 - p_{i,\bullet}) - e_i p_{i,\bullet}\,. \tag{A-16}$$

We also have

$$p_{i,0} = \int_0^X \mu_x p_{i,x} \mathrm{d}x + p_{i,X}\,. \tag{A-17}$$

and

$$p_{\bullet,0} = \int_0^X \mu_x p_{\bullet,x} \mathrm{d}x + p_{\bullet,X}\,.$$

Let us check that $\check{p}_{i,x} = p_{i,\bullet} p_{\bullet,x}$ is solution of Eq. A-1 at steady-state (i.e. Eq. A-2) with initial condition A-17. At $x = 0$,

$$\check{p}_{i,0} = p_{i,\bullet} p_{\bullet,0} = \int_0^X \mu_x p_{i,\bullet} p_{\bullet,x} \mathrm{d}x + p_{i,\bullet} p_{\bullet,X} = \int_0^X \mu_x \check{p}_{i,x} \mathrm{d}x + \check{p}_{i,X}\,,$$

which is consistent with Eq. A-17. We also have

$$\frac{\mathrm{d}\check{p}_{i,x}}{\mathrm{d}x} = -\mu_x p_{i,\bullet} p_{\bullet,x} = -\mu_x \check{p}_{i,x}\,.$$

Since

$$-e_i \check{p}_{i,x} + (c_i p_{i,\bullet} + m_i)(p_{\bullet,x} - \check{p}_{i,x}) = p_{\bullet,x}\left(-e_i p_{i,\bullet} + (c_i p_{i,\bullet} + m_i)(1 - p_{i,\bullet})\right) = 0\,,$$

(see Eq. A-16), one can equally write

$$\frac{\mathrm{d}\check{p}_{i,x}}{\mathrm{d}x} = -\mu_x \check{p}_{i,x} - e_i \check{p}_{i,x} + (c_i p_{i,\bullet} + m_i)(p_{\bullet,x} - \check{p}_{i,x})\,,$$

which is consistent with Eq. A-2.

Therefore, $p_{i,x} = p_{i,\bullet} p_{\bullet,x}$, meaning that patch age and occupancy are independent random variables for immune species.

As a consequence, $p_{i|x} = p_{i,x}/p_{\bullet,x} = p_{i,\bullet}$, hence $\pi_{i/x} = p_{i|x}/p_{i,\bullet} = 1$ for all $x$. In other words, immune species have the same relative distribution profile as infinitely fast non-immune species. The predictions made for very fast non-immune species hold for immune species as well. Pairing an immune species with any other type of species is expected to generate spurious competition.
