## Supporting Information B for "Metacommunity dynamics and the detection of species associations in co-occurrence analyses: why patch disturbance matters"

### B. Stochastic simulations

**Vincent Calcagno, Nik J. Cunniffe, Frédéric M. Hamelin**

#### B1 Monte Carlo algorithm

We first present the Monte Carlo algorithm used to sample random co-occurrence matrices. The goal is to generate a $s \times N$ co-occurrence matrix, with co-occurrence patterns drawn randomly from the steady state probabilities of the general meta-community model (Eq. A-1 in Supporting Information A).

One approach would be to compute the steady state co-occurrence probabilities for every type of patch, i.e. for every possible combination of species, similar to what is done with $q_{i,j,\bullet}$ for species pairs in Supporting Information A, section A4. Then, one can draw randomly $N$ patches from the corresponding multinomial distribution, and fill the co-occurrence matrix accordingly. However, as the number of species combinations quickly explodes with $s$, this approach is very inefficient numerically.

An efficient approach consists in the following Monte Carlo algorithm:

1. Compute the steady-state patch age distribution (Supporting Information A, section A1.2) and the steady state overall occupancies (Supporting Information A, section A1.4) once and for all;

2. Draw one random patch age $x$ from the steady-state patch age distribution (Eq. A-3 in Supporting Information A);

3. For every species, compute its steady-state probability of occupancy conditional on patch age $x$, $p_{i|x}$, from Eq. A-4 in Supporting Information A;

4. For each species, draw a random number to determine if it is present in the focal patch, based on $p_{i|x}$;

5. Complete the column of the co-occurrence matrix accordingly;

6. Repeat 2-5 for as many patches (matrix columns) as desired.

**Note:** Species that are immune to patch disturbances, if any, should have $p_{i|x}$ in Step 3 replaced with their steady state overall occupancy with $\mu_x = 0$ for all $x$ (see Supporting Information A, section A4). The latter can be computed from the classical metapopulation model results (Eq. A-11 in Supporting Information A).

The above approach is implemented as a set of R functions available at `https://github.com/nikcunniffe/MetacommunityDynamics`.

Step 2 is implemented by sampling patch ages from the distribution $p_{\bullet,x}$ defined in Eq. A-3 (in Supporting Information A) using inversion (Devroye, 1986). The requisite calculations can be done more quickly if the form adopted for $\mu_x$, the death rate for patches of age $x$, allows the user to specify

$$M(x) = \int_0^x \mu_y \mathrm{d}y$$

in closed form (this can be done by passing the argument `integratedDeathRateFunc` to the function `initPatchDeath()`). However, if an explicit functional form for $M(x)$ is not available, the values of the function at a large number of values of $x$ are pre-calculated during the initiation step of our algorithm by way of numerical integration via the library `cubature` (Narasimhan et al., 2020). This allows values of $M(x)$ for arbitrary $x$ to be obtained efficiently via linear interpolation between two pre-cached values. In either case, given a function that returns $M(x)$ for any value of $x$, the probability density of patch ages in Eq. A-3 (in Supporting Information A) can be written as

$$p_{\bullet,x} = p_{\bullet,0} \exp\left(-M(x)\right),$$

in which $p_{\bullet,0}$ is the normalisation factor ensuring $p_{\bullet,x}$ is a probability density function.

The cumulative density function for patch ages is then

$$C(x) = p_{\bullet,0} \int_0^x \exp(-M(z))\, dz.$$

Patch ages can then be sampled by choosing a uniformly distributed random number, $R$, between 0 and 1, and solving the equation $R = C(a)$ to sample the age of a single patch, $a$. This is done using the base R function `uniroot()`, with the numerical integration that is necessary to find $C(x)$ done using `cubature` (Narasimhan et al., 2020).

#### B2 Parameter values

Our illustrative set in the main article was $\mu_x = 0.2$ and $X = 20$. There was no external immigration ($m_i = 0$). The 31 species used had colonisation rates ($c_i$) in the range in $(0.33, 7.1)$ and extinction rates ($e_i$) in the range $(1e^{-3}, 3.4)$. Their overall steady-state occupancies were drawn randomly in the range $(0.3, 0.7)$. Co-occurrence matrices were sampled for $N = 1000$ sites (patches).

All details can be found in the provided R markdown file.

#### B3 Direct independence tests

We used function *fisher.test* in R to test for independence of every species pair.

All the code used can be found in the R markdown available at `https://github.com/nikcunniffe/MetacommunityDynamics`.

#### B4 Null permutation schemes

We used permutation algorithms to tests whether the partial C-score values for species pairs significantly differed from the null expectation. To this end, we used the functions provided in the R *ecosimR* package (Gotelli, 2000). We used the fixed-equiprobable (Sim2) and fixed-fixed (Sim9) permutation algorithms.

All the code used can be found in the R markdown available at `https://github.com/nikcunniffe/MetacommunityDynamics`.

#### B5 Hmsc models

We used the Hmsc package in R (Ovaskainen and Abrego, 2020) to fit joint species distribution models to null sample matrices generated from the metacommunity model. We used four different models, differing in the fixed and random effects they incorporate to describe the presence/absence probabilities. MCMC convergence, model predictions and species associations were analyzed following Tikhonov et al. (2020). The different models are summarized in Table B1.

Table B1: Hmsc models used.

| Hmsc model | Habitat covariates | Random factors† |
|---|---|---|
| M0 | none | none |
| M1 | none | 1 (levels of patch richness $1 \ldots s$) |
| M2 | 1 (log(patch age)) | none |
| M2' | 1 (patch richness) | none |

† in all models, a dummy random factor, with patch identity as a grouping factor, was also incorporated to permit the computation of species associations.

All the code used can be found in the R markdown available at `https://github.com/nikcunniffe/MetacommunityDynamics`. Note that results can vary slightly, in quantitative terms, depending on the exact sample matrix (i.e. random seed) used. The results presented were obtained on a matrix generated with 1 as a random seed, but results are representative of what one obtains with other seeds.
