## Supplementary figures and images for "Metacommunity dynamics and the detection of species associations in co-occurrence analyses: why patch disturbance matters"

### Supporting Figure 1

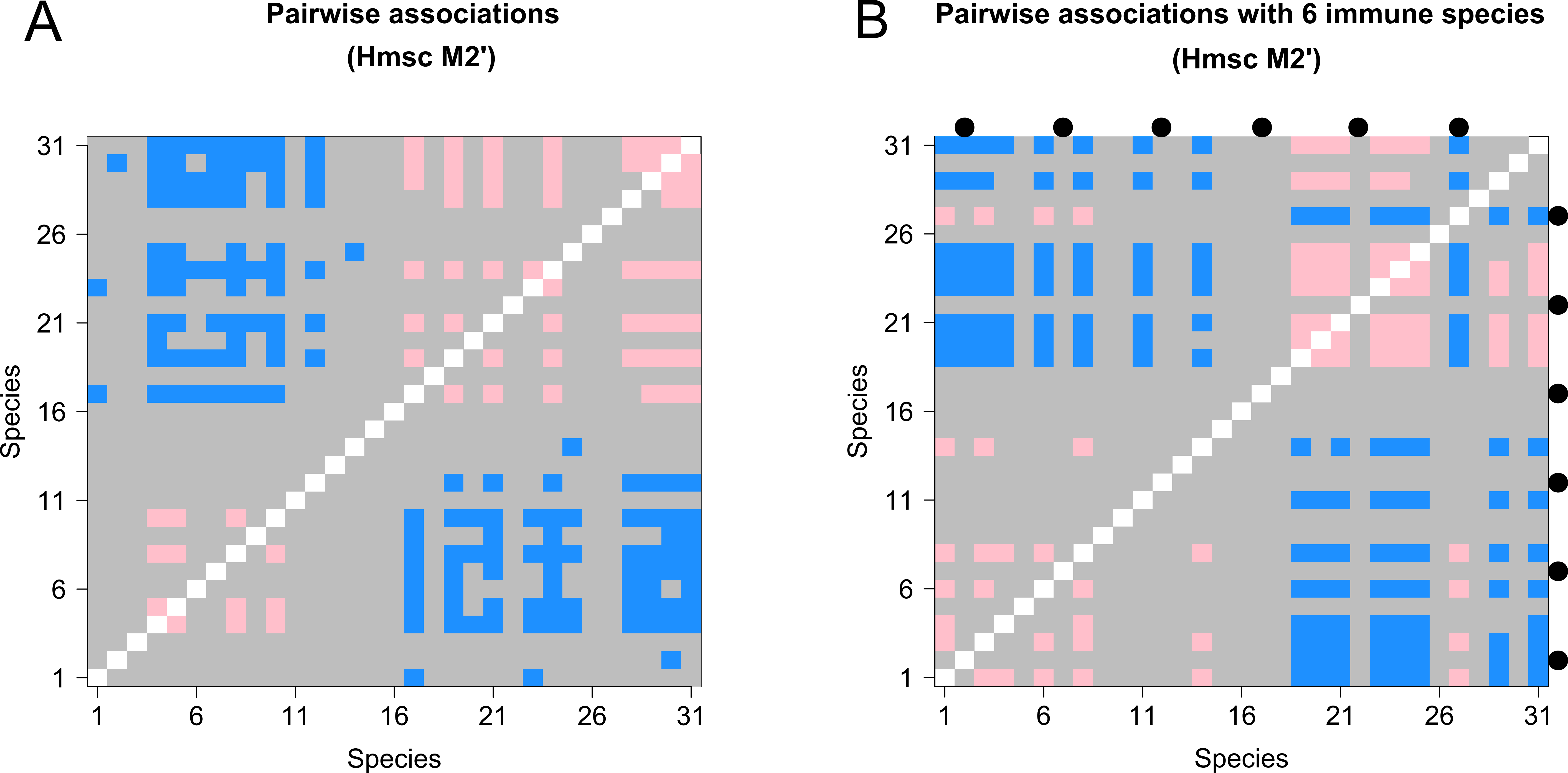
